# Profiling human hypothalamic neurons reveals a candidate combination drug therapy for weight loss

**DOI:** 10.1101/2023.07.18.549357

**Authors:** Hsiao-Jou Cortina Chen, Andrian Yang, Simone Mazzaferro, Iman Mali, Olivier Cahn, Katherine Kentistou, Christine Rowley, Natasha Stewart, Jun Wen Eugene Seah, Venkat Pisupati, Peter Kirwan, Sanya Aggarwal, Takafumi Toyohara, Mary H.C. Florido, Chad A. Cowan, Lena Quambusch, Marko Hyvönen, Matthew R. Livesey, John R.B. Perry, John C. Marioni, Florian T. Merkle

**Author notes:** These authors contributed equally (Hsiao-Jou Cortina Chen, Andrian Yang). Address for correspondence (J.C.M); (F.T.M).

## Abstract

Obesity substantially increases the risk of type 2 diabetes, cardiovascular disease, and other diseases, making it a leading preventable cause of death in developed countries. It has a strong genetic basis, with obesity-associated genetic variants preferentially acting in the brain. This includes the hypothalamic pro-opiomelanocortin (POMC) neurons that inhibit food intake and are stimulated by drugs that agonise glucagon-like 1 peptide receptor (GLP1R) including Semaglutide (Ozempic/Wegovy). We therefore hypothesised that drugs which selectively activate human POMC neurons would suppress appetite and promote weight loss, and that focusing on drugs already approved for use would facilitate rapid clinical translation. We therefore generated POMC neurons from human pluripotent stem cells (hPSCs) and identified enriched genes that were genetically associated with obesity and targeted by approved drugs. We found that human POMC neurons are enriched in GLP1R, reliably activated by Semaglutide, and their responses are further increased by co-administration of Ceritinib, an FDA-approved drug potently and selectively inhibiting anaplastic lymphoma kinase (ALK). Ceritinib reduced food intake and body weight in obese but not lean mice, and upregulated the expression of GLP1R in the mouse hypothalamus and hPSC-derived human hypothalamic neurons. These studies reveal a new potential therapeutic strategy for reducing food intake and body weight, and demonstrate the utility of hPSC-derived hypothalamic neurons for drug discovery.

## Introduction

Obesity affects billions of people worldwide, contributes to millions of excess deaths each year, and requires more effective treatment (Nyberg et al. 2018). Human genetic studies and animal models have revealed that obesity is heavily influenced by the brain, particularly in cell types involved in appetite regulation. Specifically, neurons in the arcuate nucleus (ARC) of the hypothalamus that produce proopiomelanocortin (POMC) are critical for normal body weight regulation and suppress appetite by releasing melanocyte-stimulating hormone (MSH) which binds to the downstream melanocortin 4 receptor (MC4R) (Yeo et al. 2021). Both humans and animals lacking *POMC* or *MC4R* develop hyperphagia and obesity (Y. S. Lee et al. 2006; Challis et al. 2004; Yeo et al. 2003; Ste Marie et al. 2000) whereas administering MSH or specifically stimulating POMC neurons using optogenetic or chemogenetic methods reduces food intake and promotes weight loss (Üner et al. 2019; Mountjoy et al. 2018; Zhan et al. 2013; Aponte, Atasoy, and Sternson 2011; McMinn et al. 2000).

The activity of the POMC neurons is influenced by signals such as the adipocyte-derived hormone leptin and the gut- and brain-derived hormone glucagon-like peptide-1 (GLP-1), which affect food intake, energy expenditure, and glucose metabolism through their corresponding receptors (Jais and Brüning 2022). Drugs approved for use in humans that selectively activate the GLP-1 receptor (GLP1R), including Liraglutide and Semaglutide (trade names Ozempic/Wegovy), lower blood glucose levels and promote weight loss in both animal models and humans (F. Sun et al. 2015; Gabery et al. 2020). Although the precise human cellular targets and action of GLP1R agonist remain to be fully elucidated, Liraglutide binds to and activates mouse POMC neurons (Secher et al. 2014). We therefore hypothesised that drugs which selectively activate human POMC neurons would suppress appetite and promote weight loss, and that focusing on drugs already approved for use would facilitate rapid clinical translation. Furthermore, we hypothesised that drugs targeting POMC neurons might have greater efficacy when combined with existing GLP1R agonist drugs.

It is challenging to identify such candidate drugs from the existing literature, since only a limited number have been experimentally tested for their action on POMC neurons in animal models. While these hypothesis-driven studies are clearly valuable, their throughput is low, limiting the potential for discovering novel druggable targets that might preferentially activate POMC neurons. A broader and unbiased approach is to identify genes that are enriched in POMC neurons and might therefore selectively promote POMC neuron activation when pharmacologically targeted. While single-cell and single-nucleus RNA sequencing (scRNASeq, snRNAseq) studies of mouse hypothalamic neurons permit the discovery of mouse POMC-enriched genes (Steuernagel et al. 2022), the translation of such findings to humans is notoriously poor due to species-specific functional differences (Hodge et al. 2019). To facilitate clinical translation, gene enrichment studies should ideally be performed in human POMC neurons. Several groups have published snRNAseq datasets containing human POMC neurons (W.-K. Huang et al. 2021; Herb et al. 2022; Siletti et al. 2022), but in aggregate these contain just 183 annotated POMC neurons and a modest number of genes per cell (<100), making them unsuitable for robust differential expression analysis.

To address this challenge, we improved upon existing protocols (L. Wang et al. 2015; Merkle et al. 2015) to differentiate human pluripotent stem cells (hPSCs), into hypothalamic neurons, including POMC neurons. These hPSC-derived POMC (hPOMC) neurons can be generated at scale and have revealed human-specific aspects of neuropeptide processing and obesity-associated cellular phenotypes (Kirwan et al. 2018; L. Wang et al. 2019; Rajamani et al. 2018). Using scRNAseq, we first established that hPOMC neurons are transcriptionally similar to their counterparts in the human neonatal hypothalamus (W.-K. Huang et al. 2021). We then performed genetic association analysis using a recently described variant to gene mapping approach that leverages large-scale population-based datasets to identify enriched genes that are associated with childhood and adult human obesity (Kentistou et al. 2023). Among the candidate genes, we prioritised those that had drugs approved for use in humans that were likely to cross the blood-brain-barrier for testing in hPOMC neurons and in mouse models of diet-induced obesity (DIO) to enable their potentially rapid clinical translation.

Specifically, we identified and focused on anaplastic lymphoma kinase (ALK) which is predominantly expressed in the brain and has been associated with cancer and body weight regulation when aberrantly activated or functionally perturbed (Orthofer et al. 2020; Soda et al. 2007). ALK is a receptor tyrosine kinase in the insulin receptor superfamily (Morris et al. 1997) and its activation triggers the same signal transduction pathways (JAK/STAT3, PI3K/AKT, RAS/MAPK) utilised by the insulin, leptin, and GLP1R, providing opportunities for signal integration. We found that the potent, selective, and FDA-approved small molecule ALK inhibitor Ceritinib potently reduced body weight in DIO mice, but not in lean mice. Co-administration of Semaglutide resulted in even greater weight loss, and Ceritinib treatment consistently upregulated the expression of GLP1R in the mouse hypothalamus as well as in hPSC-derived hypothalamic neurons, which showed potentiated responses to Semaglutide. Together, these findings suggest potential functional interaction between the ALK and GLP1R pathways and raise the intriguing possibility of combinatorial therapeutic strategy for the treatment of human obesity. This study also establishes hPSC-derived hypothalamic neurons as a powerful platform for anti-obesity drug discovery.

## Results

### Generation of human hypothalamic neurons across multiple genetic backgrounds and culture conditions

Our aim was to identify obesity-associated genes that are differentially expressed in hPOMC-expressing neurons and are targets of known drugs which could potentially be repurposed to treat obesity (Fig. 1A). To this end, we differentiated six different hPSC lines (HUES9-POMC-GFP, KOLF2.1J, NN0003932, NN0004297, NCRM5, and PGP1, Table S1) into hypothalamic neurons as previously described (L. Wang et al. 2015; Merkle et al. 2015; H.-J. C. Chen et al. 2023; Kirwan, Jura, and Merkle 2017) (Fig. 1B, S1A). To enhance neuronal maturation and test the robustness of our findings under different environmental conditions, we cultured them in four different maturation conditions including standard N2B27-based medium alone or with primary mouse cortical astrocytes (NB, NBA), as well as Synaptojuice medium alone or with astrocytes (SJ, SJA) (Fig. 1B). To enable hPSC-POMC neurons to be functionally studied, we used CRISPR/Cas9 genome editing to insert the T2A ribosomal skipping sequence followed by a green fluorescent protein (GFP) into the stop codon of the endogenous human *POMC* gene in the HUES9 embryonic stem cell line to create a knock-in reporter that faithfully expresses GFP in POMC neurons (Fig. 1C, S1B-G). Upon immunostaining these hPSC-derived hypothalamic cultures on Day 35 ± 2, we observed cells with neuronal morphology that were strongly immunopositive for a POMC-specific antigen and co-expressed the neuron-enriched gene microtubule-associated protein 2 (MAP2) (Fig. 1D). In all tested cell lines, we found that the percentage of MAP2 immunopositive cells that were also immunopositive for POMC was significantly higher in both the SJ and SJA conditions than in NB (Fig. 1E). Together, we found that hPOMC neurons were generated across all cell lines and culture conditions, in some cases at up to 45% efficiency (Fig. 1E). At the time of analysis, GFAP immunopositive cells only made up a small fraction of all culture cells (Fig. S1H), and we did not observe significant effects of astrocyte co-culture on the fraction of POMC immunopositive cells.

**Figure 1.**
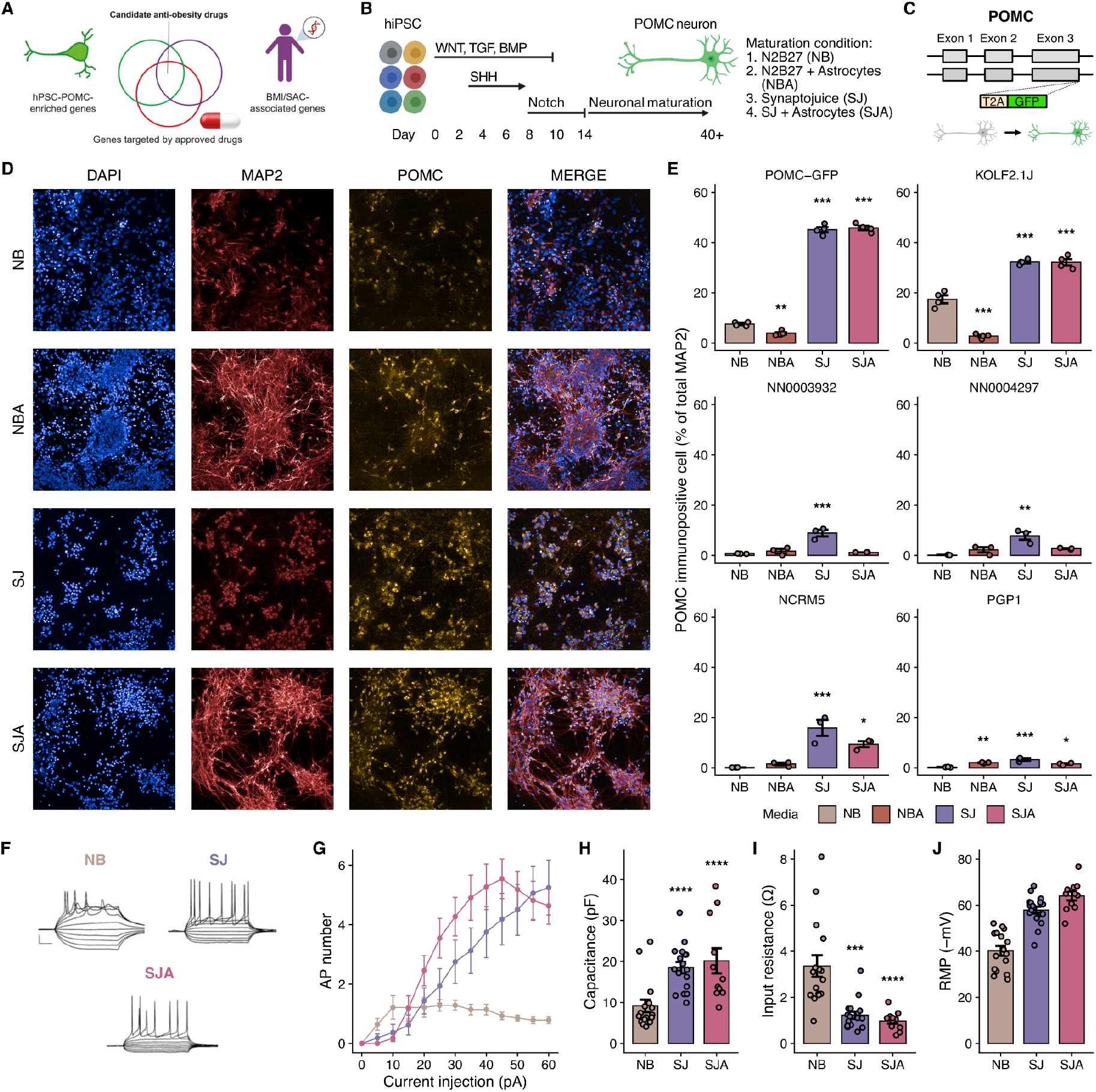
Generation of hypothalamic neurons from human pluripotent stem cells (hPSCs) across multiple genetic backgrounds and culture conditions. **A)** Schematic of project overview to identify obesity-associated genes that are enriched in hPSC-derived proopiomelanocortin (POMC) neurons and are targets of drugs approved for human use. **B)** Experimental schematic of human hypothalamic differentiation and maturation in different media including N2B27 (NB), N2B27 + astrocytes (NBA), Synaptojuice (SJ), and Synaptojuice + astrocytes (SJA). **C)** Schematic diagram of an endogenous knock-in *POMC-GFP* reporter cell line. **D)** Representative images of HUES9 *POMC-GFP*-derived hypothalamic cultures stained with DAPI (blue), MAP2 (red), and POMC (yellow) across the 4 different culture conditions, NB, NBA, SJ, and SJA. **E)** The quantified percentage of MAP2^+^ cells expressing POMC is displayed for six different hPSCs lines used in this study including HUES9-POMC-GFP, KOLF2.1J, NN0003932, NN0004297, NCRM5, and PGP1 (n=3-4). **F)** Representative whole-cell current-clamp recording traces of single POMC-GFP^+^ neurons maintained in NB, SJ, and SJA. Traces depict evoked potential activity in response to depolarizing and hyperpolarizing 500 ms current injection. Scale bar, 20 mV, 50 ms. **G)** Action potential output of POMC-GFP^+^ neurons maintained in NB (n = 14, N = 3), SJ (n = 16, N = 3), and SJA (n = 11, N = 3) in response to depolarising current injection. **H-J)** Intrinsic membrane properties of the POMC-GFP^+^ neurons including whole-cell capacitance, input resistance, and resting membrane potential. Data represent mean ± SEM; *p<0.05, **p<0.01, ***p<0.001.

Since the intrinsic excitability of POMC neurons changes over the course of their maturation (Roberts et al. 2019), we used patch-clamp electrophysiology to study the intrinsic membrane properties of GFP^+^ hPOMC neurons to assess their maturation level. Excitability was assessed by the ability of GFP^+^ hPOMC neurons to fire action potentials in response to depolarising current injection. Cells cultured in SJ and SJA fired more action potentials in response to current injection (Fig. 1F & 1G), as expected from functionally more matured neurons. Furthermore, SJ and SJA neurons displayed more hyperpolarised resting membrane potentials, decreased input resistance (p<0.001) and increased whole cell capacitance (p<0.001; Fig. 1H-J) in comparison to NB-cultured neurons, which broadly correspond to values observed in postnatal POMC neurons (Roberts et al. 2019). These data demonstrate that SJ and SJA cultures are electrophysiologically more mature, which together with the increased efficiency of hPOMC neuron generation, prompted us to select these conditions for further study.

### Single-cell characterisation of in vitro-generated hypothalamic cultures

Since hypothalamic directed differentiation from hPSCs gives rise to heterogeneous cell populations (W.-K. Huang et al. 2021; Merkle et al. 2015), we sequenced single cells to catalogue the cell types present in 40 ± 2 day old cultures produced from six cell lines and four growth conditions (Fig. S2A). After quality control, genetic demultiplexing, and doublet removal, we retained 152,356 high-quality cells from 17 distinct single-cell cDNA libraries (Fig. S2B). We next analysed all cells and found that cell line PGP1 acted as an outlier by generating relatively few neurons, so we removed it from the dataset (Fig. S2C-E). After Louvain clustering and further quality control of the resulting dataset, we found that all cell lines contributed to similar cell clusters (Fig. S2F). Patterns of gene expression were consistent with the predominantly ventral and anterior hypothalamic identity of hPSC-derived cells, as suggested by the expression (UMI > 1) of the transcription factor (TF) *NKX2.1* and/or *SIX3* in a majority (124,046/142,099, 87.30%) of hPSC-derived cells, less abundant expression of the telencephalon-enriched TF *FOXG1* (16,475/142,099, 11.59%), and rare expression of *FOXB1, IRX3,* or *PITX2* (4,172/142,099, 2.94%) that are enriched in the caudal hypothalamus and more posterior brain structures. We observed a higher proportion of neuron generation in cells matured in SJ than in NB based on cell clusters that expressed high levels of *STMN2* and *TUBB3 (TUJ1)* and relatively low levels of nestin (*NES*) (Fig. 2A-D). We further found a higher proportion of neurons in cultures matured in SJ than in NB, whereas the effects of astrocyte co-culture were modest (Fig. S2G). We therefore separated our single-cell data based on the maturation media used (e.g. NB and NBA or SJ and SJA) and re-analysed them separately to better characterise the hypothalamic cell types generated (Fig. 2).

**Figure 2.**
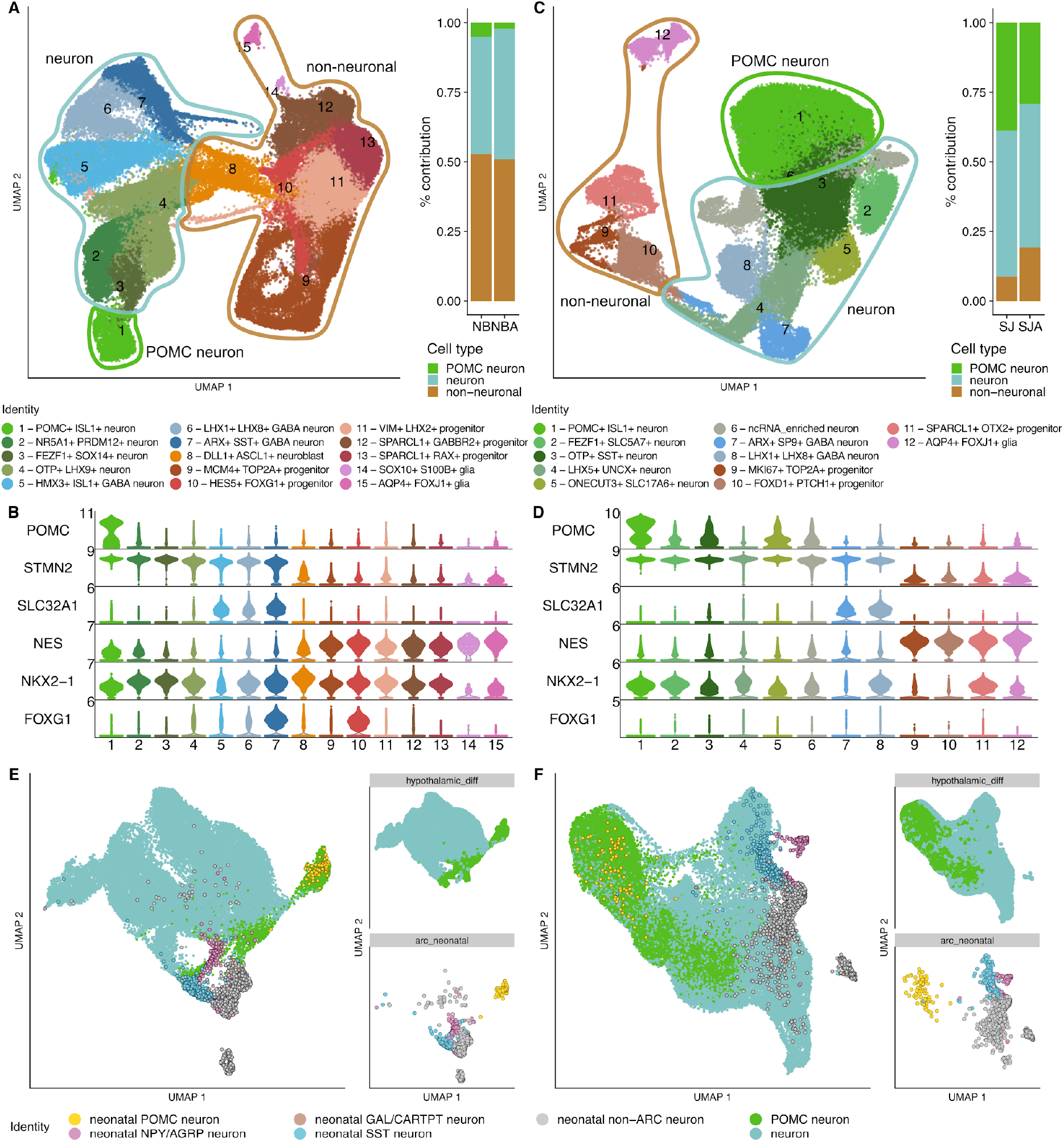
Transcriptional identity of hPSC-derived hypothalamic neurons. **A,C)** UMAPs of scRNAseq data from hPSCs differentiated to hypothalamic cells reveals similar cell populations regardless of whether they are matured in N2B27-based medium (NB, NBA; A), or SJ-based medium (SJ, SJA; C) in which the overall efficiency of neuron (cyan) and POMC neuron (green) production is higher (insets). **B,D)** Broadly, gene expression patterns define cell populations as neurons (shades of green), GABAergic neurons (shades of blue), or progenitors and glia (shades of red and brown). **E,F)** Integration of scRNAseq data from hPSC-derived cultures differentiated in N2B27-based medium (E), or SJ-based medium (F) with single nucleus RNAseq data from the neonatal human hypothalamus reveals the transcriptional similarly of primary human POMC neurons (yellow) and hPSC-derived POMC neurons (green).

### Molecular identity of other hPSC-derived cell clusters

To gain insight into the identity of cells within each cluster, we used a combination of unbiased differential gene expression between clusters and manual annotation with genes indicative of hypothalamic development and cell identity. Our analysis revealed 15 clusters of distinct cell types cultured in NB and 12 such clusters in the SJ matured cells (Fig. 2A-D, Fig. S3A-B), which had likely counterparts *in vivo* based on their expression of regionally enriched TFs and/or genes enriched in radial glia, tanycytes, or ependymal cells (Kim et al. 2020; Bedont, Newman, and Blackshaw 2015; Alvarez-Bolado, Grinevich, and Puelles 2015; Ma et al. 2021). In the NB condition, seven cell clusters were neuronal and three were likely GABAergic given their expression of the GABA transporter *SLC32A1* and *DLX2* (Fig. 2A, green and blue shading, respectively) (Lindtner et al. 2019). One cell cluster was highly enriched for *POMC* and *ISL1,* suggesting a POMC neuron and hypothalamic arcuate nucleus (ARC) identity (B. Lee et al. 2016) (Fig. 2A,B, S3A). A second cluster was enriched in *NR5A1(SF1)* and *PRDM12* indicative of an ARC and ventromedial hypothalamus (VMH) identity, as was a third VMH-like cell cluster enriched in *FEZF1* and *SOX14 (Kurrasch et al. 2007)*. Cells from a fourth cluster were enriched in *OTP*, *SST* and *LHX9* suggesting an identity of the ventrolateral anterior hypothalamus (vlAH) and/or intra-hypothalamic diagonal (ID) (Shimogori et al. 2010). Among the GABAergic neuron clusters, the fifth cluster was enriched in *HMX2/3*, *GSX1,* and *ISL1* which are co-expressed in the preoptic area (POA) (Gelman et al. 2009; W. Wang et al. 2004), and the sixth was enriched in *LHX1*, *LHX8*, *ARX*, and *UNCX* suggesting a suprachiasmatic nucleus (SCN)/dorsolateral anterior hypothalamus (dlAH), and/or ID identity (Shimogori et al. 2010; Bedont et al. 2014), and the seventh cluster was enriched in *FOXG1*, *DLX2*, and *SST* and likely corresponded to the medial and lateral ganglionic eminences (MGE/LGE) of the ventral forebrain (Hu et al. 2017). The eighth cluster contained *ASCL1*-enriched neuroblast-like cells (Guillemot et al. 1993), and all remaining clusters were enriched in *NES* and *VIM*, indicating their likely progenitor or glial identity (Guillemot et al. 1993; Lendahl, Zimmerman, and McKay 1990). Indeed, one cluster was highly enriched for the proliferation-associated genes *MKI67* and *TOP2A*, suggesting ongoing cell division among these radial glial-like progenitors. Other non-neuronal clusters were enriched in markers associated with different tanycyte populations including *HES5* and *FRZB* (β-tanycyte), *LHX2* and *FOXD1* (ventral-anterior hypothalamus (vAH) tanycyte-like), *SPARCL1* (tanycyte-like), *RAX*, *SLCO1C1*, and *COL25A1* (β-tanycyte) (R. Chen et al. 2017; Newman et al. 2018; Miranda-Angulo et al. 2014; Prevot et al. 2018; Sullivan, Potthoff, and Flippo 2022). Finally, one small cell cluster appeared to consist of oligodendrocyte-like cells enriched in *S100B*, *FOXD3*, *SOX10*, and *MPZ (Elmentaite et al. 2021)*, whereas cells in the final cluster resembled ependymal cells due to the enrichment in *FOXA1*, *FOXJ1*, and *AQP4* but relative absence of *GFAP* (MacDonald et al. 2021; Czerny et al. 2022).

SJ-matured cultures contained many similar clusters (Fig. 2C,D, S3B), but had a higher percentage of neurons distributed among seven cell clusters (Fig. 2A,C), of which one was again highly enriched in *POMC* and *ISL1* corresponding to POMC neurons of the ARC (Fig. 2D). Based on patterns of enriched gene expression as described above, the remaining neuronal clusters likely correspond to the VMH (*SOX14* and *FEZF1*), the vlAH (*OTP* and *SST*, along with *ARX* and *NKX2.2*), the SCN and dorsolateral anterior hypothalamus (dlAH), (*LHX1* and *LHX8,* along with *LHX5, LHX9*, *SIM1*, and *UNCX*), and a *ONECUT3* and *SLC17A6* enriched cell population that has been mapped to paraventricular nucleus (PVN) of the dlVH, as well as to the lateral hypothalamus, zona incerta, and the mediolateral zone (Zupančič et al. 2023). Although under SJ conditions there were fewer cycling cells than in NB, some *MKI67* and *TOP2A* expressing cells persisted, as did radial glial-like cells of the vAH (*FOXD1, NKX2-2, ASCL1,* and *DLL1*), tanycyte-like cells of the vAH (*SPARC*, *HES5, OTX2,* and *RAX*), and a distinct population of ependymal-like cells (*FOXA1, FOXJ1,* and *AQP4*) (Fig. 2C,D, S3B). Together, these data suggest that our differentiation protocol predominantly generated anterior and ventral hypothalamic nuclei such as the ARC, VMH, and adjacent structures.

### hPSC-derived POMC neurons resemble their counterparts in the human brain

To determine how closely hPOMC neurons resemble their counterparts *in vivo*, we compared single-cell transcriptomes of neurons matured in either NB or SJ media to single-nucleus transcriptomes of human neonatal ARC neurons including primary POMC neurons (neonatal pPOMC) (W.-K. Huang et al. 2021). We found that these datasets integrated readily and that neonatal pPOMC neurons clearly overlapped with the predominant cluster of hPOMC neurons regardless of their genetic background or the culture conditions in which they were grown (Fig. 2E,F). Other primary neonatal ARC cell types also overlapped to some extent with hPSC-derived hypothalamic cells, including neonatal SST and AGRP/NPY neurons. To quantify and confirm these observations, we transferred labels from primary ARC neurons onto hPSC-derived hypothalamic neurons and found that hPOMC neurons were predominantly assigned the primary neonatal ‘POMC’ label (Fig. S3C,D). Together, these results suggest that hPOMC neurons generated *in vitro* have a similar molecular identity to *bona fide* human POMC neurons.

### Identification of druggable candidate genes targeting human POMC neurons

To identify genes enriched in POMC neurons that might facilitate their preferential activation, we analysed hPSC-derived hypothalamic cultures matured in SJ and found they contained 22,892 hPOMC neurons, each containing an average of ∼22,714 unique molecular identifiers (UMIs) and ∼5,827 genes per cell (Fig. 2C, Fig. S3B). These figures compare favourably to the ∼449 UMIs and ∼79 genes per cell seen across 183 annotated POMC neurons in three published studies with primary human samples (W.-K. Huang et al. 2021; Herb et al. 2022; Siletti et al. 2022). We therefore split our dataset into hPOMC or non-POMC neurons, and pseudo-bulked the cells within each of these groups for each cell line or experimental replicate for differential expression analysis. Overall, we compared 18,921 genes expressed in the 22,892 hPOMC neurons to those expressed in the 35,643 non-POMC human neurons derived in parallel from the same cultures, and identified 575 genes that were significantly (log2(fold-change) > 0.5 and FDR < 0.05) enriched in hPOMC neurons, including genes such as *GLP1R* with known relevance to body weight regulation. (Fig. 3A, Table S2A).

**Figure 3.**
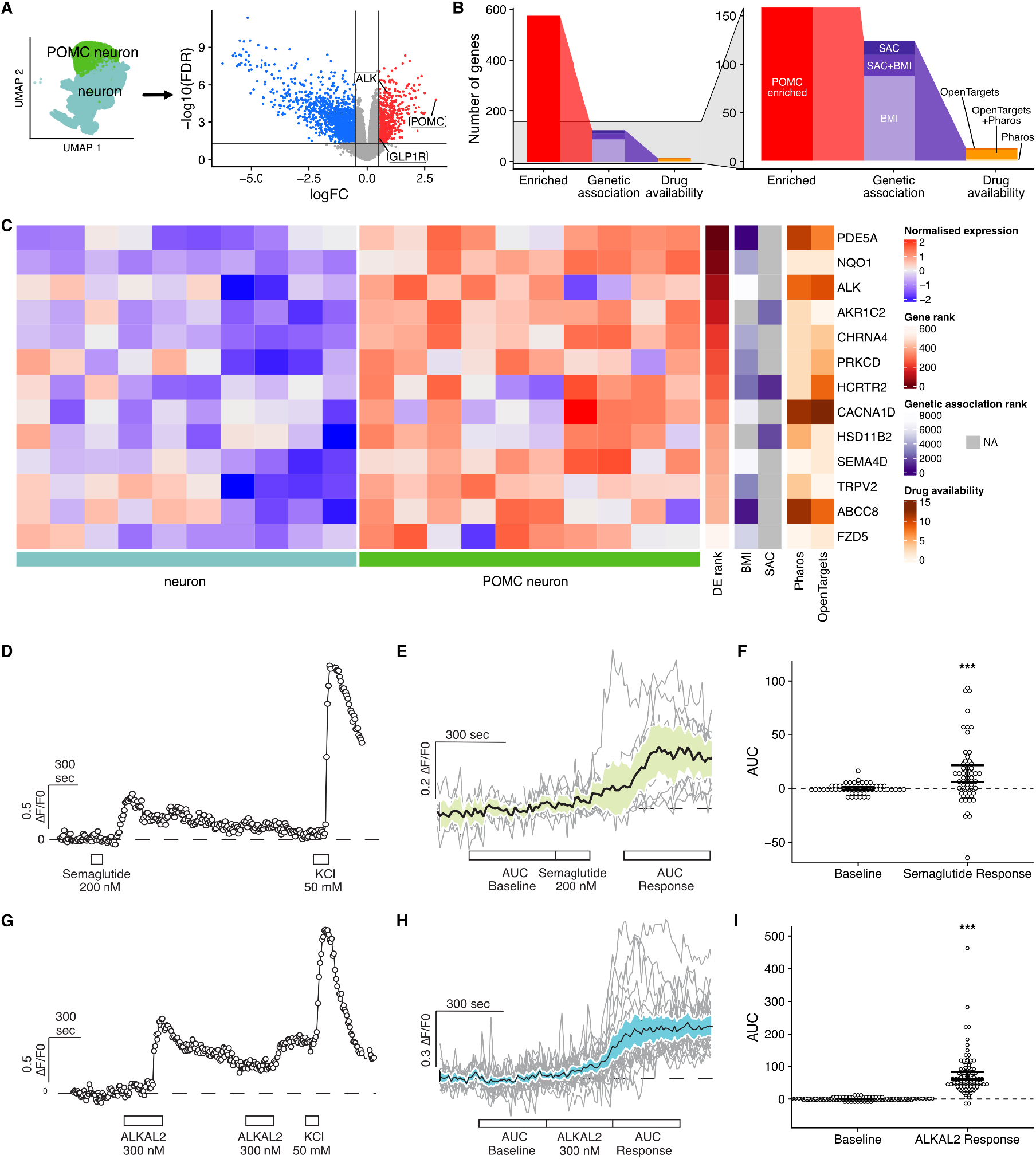
Identification of candidate drugs to repurpose for body weight loss. **A)** Single-cell RNAseq data from hPSCs differentiated to hypothalamic neurons and matured in SJ-based media were restricted to ‘neuron’ and ‘POMC neuron’ populations (left UMAP plot, cyan and green, respectively) that were pseudo-bulked for differential gene expression analysis for each sample to reveal neurons enriched in human POMC neurons (right volcano plot, red dots). **B)** Schematic of how POMC enriched genes (red) were prioritised based on their genetic association (purple) with with adult body mass index (BMI) or body size at childhood (SAC) and finally by the availability of drugs approved for use in humans as represented in the Pharos or OpenTargets databases (orange). **C)** Z score-normalised gene expression (blue to red colour scale) of genes ranked by their enrichment in POMC neurons (shades of red), their genetic association with human obesity (shades of purple) and the number of available drugs (shades of orange) revealed 25 genetic targets with candidate drugs for repurposing, including glucagon-like peptide 1 receptor (*GLP1R*) and anaplastic lymphoma kinase (*ALK*) (see volcano plot in A). **D,E)** Calcium imaging traces of human hypothalamic neuron responses to Semaglutide as illustrated by a representative trace (D) and population average (E, n=57, N=3). **F)** Semaglutide induced a significantly higher mean area under the curve (AUC). **G,H)** ALKAL2, a peptide agonist of the ALK receptor, robustly increased intracellular calcium concentrations in human hypothalamic neurons: representative trace (G), population average (H, n=109, N=2), **I)** Mean AUC is significantly higher upon ALKAL2 administration. Data represent mean with 95% CI; *p<0.05, **p<0.01, ***p<0.001.

Given the central role that POMC neurons play in body weight regulation, we next asked which of our identified hPOMC-enriched genes have human genetic support for a direct involvement in obesity. Specifically, we employed a recently developed ‘GWAS to Genes’ pipeline (Kentistou et al. 2023) that prioritises candidate causal genes proximal to GWAS association signals. We ran this on a GWAS meta-analysis (up to N=806,834) for adult BMI (Yengo et al. 2018), in addition to a measure of recalled childhood body size (N=444,345) in UK Biobank (Bycroft et al. 2018) which has previously been validated against objectively measured childhood BMI (Felix et al. 2016). Using these data, we found that 109 and 36 of the differentially expressed genes were proximal (±500 kb) to genome-wide significant signals for adult BMI and childhood body size, respectively (Table S2C). Many of these associated genetic variants were linked to candidate genes through multiple lines of evidence. For example, *ADAM23* and *TRIM47* are both the nearest genes to variants associated with adult BMI, which reside within enhancers and influence expression of these genes.

Next, we hypothesised that targeting some of these candidate genes by repurposing drugs approved for human use might selectively activate POMC neurons to promote weight loss and provide a rapid path to the clinic. We therefore searched the Pharos and Open Targets databases of drugs in clinical trials or already approved for use in humans by the FDA. Among the 575 genes enriched in SJ-matured hPOMC neurons 43 were present in Pharos, 49 were in Open Targets, and 13 were also genetically associated with obesity (Fig. 3C, Table S2C). Eleven of these genes (85%) replicated when we repeated the analysis with NB-matured cells (Fig. S3C, Table S2D). We then manually reviewed the literature to identify genes that were enriched in the brain and/or hypothalamus, had experimental evidence supporting their role in body weight regulation, and were targeted by drugs predicted to cross the blood-brain barrier (BBB) to directly act on their central targets. Based on our manual curation, we selected anaplastic lymphoma kinase (*ALK*), NAD(P)H dehydrogenase 1 (*NQO1*), hypocretin receptor type 2 (*HCRTR2*), and cholinergic receptor nicotinic alpha 4 subunit (*CHRNA4*) for further study.

In addition to these genes, we also prioritised *GLP1R* for functional testing *in vitro* and *in vivo* for several reasons: it is enriched in hPOMC neurons (Table S2A), genetically associated with childhood BMI (Helgeland et al. 2022), and is also the target of drugs such as Semaglutide (a.k.a. Wegovy, Ozempic, Rybelsus). These drugs mimic the incretin hormone GLP-1, and are rapidly growing in popularity for the treatment of T2D and obesity (Drucker 2018). They act in part by stimulating neuron populations that reduce food intake, including hindbrain neurons and hypothalamic POMC neurons (Secher et al. 2014; Burmeister et al. 2017). Furthermore, the combined activation of GLP1R and other receptors, such as the gastric inhibitory polypeptide receptor (GIPR) and/or the glucagon receptor (GCGR), can produce an even greater reduction in appetite and food intake (Müller et al. 2022), suggesting that they could form the basis of other effective combinatorial therapies targeting hPOMC neurons.

### Functional confirmation of GLP1R and ALK activity in hPOMC neurons

Of the genes we had identified, we were particularly interested in ALK, a receptor tyrosine kinase in the insulin receptor superfamily that is predominantly expressed in the brain (Iwahara et al. 1997; Morris et al. 1997; Vernersson et al. 2006) and is activated by the endogenous peptide ligand ALKAL2 (Bilsland et al. 2008; Guan et al. 2015; Reshetnyak et al. 2021). Its kinase domain is homologous to that of the insulin receptor and its activation leads to signal transduction via PI3K-AKT, JAK-STAT, and RAS/MAPK pathways (Palmer et al. 2009; Zamo et al. 2002). The genetic fusion of ALK with other proteins can lead to its unregulated activation to promote the emergence of non small-cell lung cancer and other cancer types (Hallberg and Palmer 2016), whereas its loss of function in flies and mice confers resistance to diet-induced obesity (Orthofer et al. 2020). ALK knockout has not been reported to adversely affect development, behaviour, or cognitive function (Bilsland et al. 2008), and may extend lifespan in flies (Woodling et al. 2020), suggesting its long-term inhibition would not be detrimental. Specifically, the drug Ceritinib (a.k.a. Zykadia) is a potent (IC_50_ 0.2 nM) and selective second-generation inhibitor of ALK’s kinase domain (Marsilje et al. 2013; Mok et al. 2017) that is able to cross the blood brain barrier (BBB) (Chow et al. 2022; Shaw et al. 2014; Kort et al. 2015)

To test whether GLP1R and ALK receptors are functional in hPSC-derived hypothalamic neurons, we exposed 40 ± 4 day-old cultures to agonists of these receptors and used calcium imaging to measure their responses. We found significantly increased (p<0.001) relative fluorescence (ΔF/F) in hPSC-hypothalamic neurons after incubation with 200 nM of Semaglutide for 2 minutes, indicative of an increased intracellular Ca^2+^ concentration (Fig. 3D-F). To stimulate ALK, we synthesised its endogenous ligand, the peptide ALK And LTK Ligand 2 (ALKAL2). After exposing hypothalamic cultures to 300 nM of purified ALKAL2 for 5 minutes, we again observed significantly increased (p<0.001) relative fluorescence in recorded cells (Fig. 3G-I). Together, these findings indicate that hPSC-derived hypothalamic neurons express both GLP1R and ALK and respond to their respective agonists.

### The ALK inhibitor drug Ceritinib reduces body weight selectively in obese mice

Since constitutive knockout mice of both ALK and and its ligand ALKAL2 (Augmentor α) exhibited similar thinness phenotype and resistance to DIO (Orthofer et al. 2020; Ahmed et al. 2022), we hypothesised that pharmacological inhibition of ALK with the potent FDA-approved drug Ceritinib would also reduce body weight in a mouse model of DIO. To test this hypothesis, we fed mice for 1 week with either a 60% high fat diet-fed (HFD) or a nutritionally and fibre-matched control diet (CD), and then intraperitoneally injected them with a single dose of vehicle (Veh; dimethyl sulfoxide, DMSO) or Ceritinib (Cer, 10 mg/kg) (Fig. 4A). This dose of Cer is similar to that given to cancer patients (∼6 mg/kg/day) (Dhillon and Clark 2014). We found that CD-fed mice showed a similar and expected increase in body weight over 24 hours regardless of the injection group they were randomly assigned to (Fig. 4B). In mice fed with HFD, we observed that Veh-treated mice also gained weight over 24 hours whereas Cer-treated mice lost 2.5 ± 0.63% of their body weight and significantly diverged (p<0.01) from the Veh group (Fig. 4B). These results suggest that Ceritinib acts specifically on energy balance, since non-specific side effects such as nausea or malaise should reduce body weight in both HFD and CD groups.

**Figure 4.**
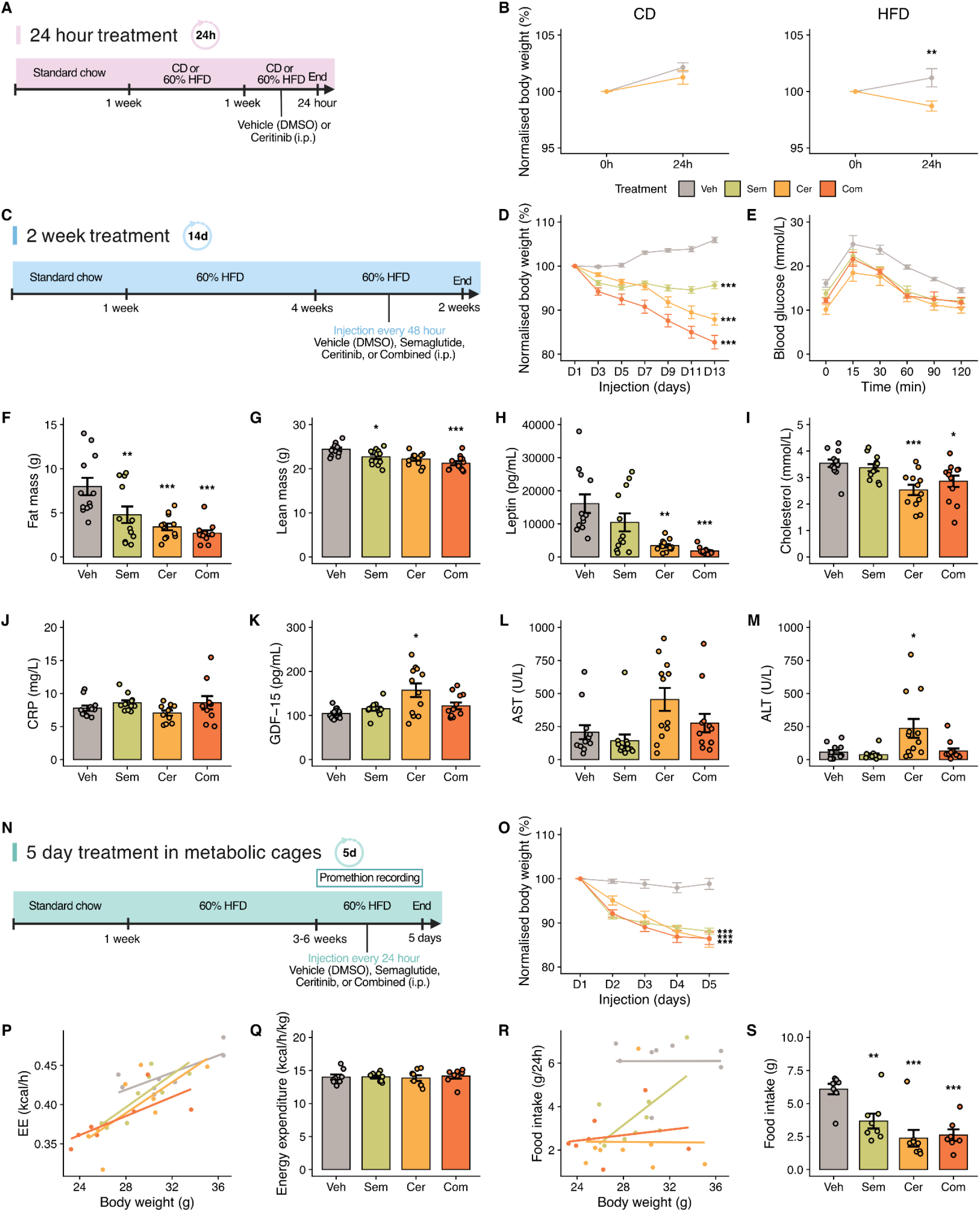
The ALK inhibitor Ceritinib significantly reduces fat mass and food intake in DIO mice. **A)** Experimental schematic to test the acute effects of vehicle (Veh; DMSO) or Ceritinib (Cer) injection in mice fed for 1 week of 60% high fat diet (HFD) or a nutritionally matched control diet (CD) with an experimental end-point 24 hours later. **B)** Mice fed CD gained weight at a similar rate regardless of injection condition (n=10/group), whereas HFD-fed mice significantly lost weight when treated with Cer (n=5/group), suggesting it acts to normalise excess energy balance rather than inducing nausea or non-specific toxicity. **C)** Experimental schematic of the long-term effects of Veh, Cer, Semaglutide (Sem) or a combination of Ceritinib and Semaglutide (Com) given every 48 hours to DIO mice. **D)** Sem, Cer, and Com treatment all induce significant body weight loss relative to Veh controls, with the greatest effect observed in the Com group (n=12/group). **E)** Weight loss is associated with a significant improvement in glucose homeostasis as revealed by an intraperitoneal glucose tolerance test (n=4/group). **F,G)** EchoMRI measurements (n=12/group) reveal that fat mass is dramatically reduced by Sem, Cer, and Com treatment (F), and lean mass is also modestly but significantly reduced in these groups (G). **H-I)** The loss in fat mass is accompanied by a significant reduction in plasma concentrations of leptin (H) and cholesterol (I) in Cer and Com groups (n=12/group). **J-M)** Plasma concentrations of c-reactive protein (CRP) were not elevated in any treatment group relative to Veh (J), but GDF-15 (K) and the liver enzymes aspartate aminotransferase (AST, L) and alanine aminotransferase (ALT, M), were moderately but significantly elevated in only the Cer group, suggesting that drug toxicity does not drive the dramatic weight loss observed with Com treatment (n=12/group). **N)** Experimental schematic of metabolic measurements in DIO mice over 5 days of treatment. **O)** Body weight recorded prior to daily injection (n=8/group) followed a similar trajectory to earlier studies (D). **P-S)** In contrast to previous reports in *Alk* deficient mice, acute Alk inhibition with Cer or Com had no effect on energy expenditure (Q-R) but significantly reduced food intake (S-T) (n=8/group). Data represent mean ± SEM; *p<0.05, **p<0.01, ***p<0.001.

### Ceritinib and Semaglutide reduce body weight and fat mass in DIO mice

To measure how extended treatment of Ceritinib affects body weight and to test for potential interactions between Ceritinib and Semaglutide, we generated DIO mice by feeding them for 4 weeks of 60% HFD, and maintained them on this diet while injecting them with candidate drugs every 48 hours for 2 weeks (Fig. 4C). We randomly allocated mice into groups and injected them with either Veh (DMSO), Cer (10 mg/kg), Semaglutide (Sem; 40 μg/kg), or a combination of Ceritinib and Semaglutide (Com). Mice in the Veh group gained 5.92 ± 0.6% in body weight at D13 compared to D1 (Fig. 4D), whereas Sem mice lost 4.34 ± 0.8% of their body weight during the same period. Strikingly, the Cer or Com groups lost 12.11 ± 1.28% or 17.28 ± 1.52% of their body weight over this period, respectively. When considering body weights measured at D13 and normalised to D1, drug treatment groups of Sem (95.67 ± 0.8%), Cer (87.90 ± 1.28%), and Com (82.72 ± 1.52%) were all significantly different (p<0.001) from the Veh group (105.92 ± 0.60%). Glucose tolerance tests (GTT) performed at the end of the 2 week treatment revealed that these reductions in body weight were accompanied by a marked improvement in glucose tolerance in all drug-treated groups at 30 (Veh vs Sem, p<0.05; Veh vs Cer, p<0.01; Veh vs Com, p<0.05), 60 (p<0.01), and 90 (Veh vs Sem, p<0.05; Veh vs Cer, p<0.01; Veh vs Com, p<0.05) minutes when compared to vehicle controls (Fig. 4E). In contrast, when we injected drugs targeting other candidate genes (Fig. 3C) including NAD(P)H quinone dehydrogenase 1 (*NQO1*), hypocretin receptor type 2 (*HCRTR2*), and cholinergic receptor nicotinic alpha 4 subunit (*CHRNA4*) in DIO mice, we did not observe a significant reduction in body weight (Fig. S5)

To test how body composition was affected by drug treatment, we performed Echo MRI and found that fat mass was significantly reduced in Sem (p<0.01), Cer (p<0.001), and Com (p<0.001) groups (Fig. 4F). Lean mass was also reduced to a smaller extent (Fig. 4G) as reported in previous studies with Semaglutide-treated DIO mice (Martins et al., 2022) or ALK knockout mice (Orthofer et al. 2020). To corroborate these studies, we measured plasma leptin concentrations after 2 weeks of drug treatment and found it was significantly reduced in the Cer (p<0.01) and Com (p<0.001) groups, likely due to decreased fat mass (Fig. 4H). Furthermore, Cer and Com treated mice had significantly reduced levels of plasma cholesterol (p<0.001 and p<0.05, respectively. Fig. 4I), consistent with reported genetic variants at the *ALK* locus are associated with plasma cholesterol concentrations (Orthofer et al. 2020). Together, these results suggest that pharmacological Alk inhibition dramatically reduces body weight and fat mass, particularly when combined with GLP1R stimulation.

### Ceritinib does not primarily act by non-specific toxicity

While our acute injection studies (Fig. 4B) suggested that Ceritinib did not induce weight loss due to acute nausea or toxicity 24 hours after injection, we wondered if long-term exposure might lead to tissue damage, toxicity, or off-target effects that might contribute to the observed weight loss. We therefore injected animals for 2 weeks and collected plasma from animals 2 hours after the final drug injection to capture both acute and chronic responses to drug administration. We first tested for changes in the circulating levels of c-reactive protein (CRP), a sensitive marker of stress and inflammation, and did not observe significant elevation of CRP in any group relative to vehicle-treated controls (Fig. 4J). Next, we tested for elevation in the circulating levels of growth differentiation factor 15 (GDF-15) which is secreted from many tissues upon activation of the cellular integrated stress response (Patel et al., 2019) and found that levels were moderately elevated (p<0.05) in mice treated with Cer, but not Com (Fig. 4K). This finding prompted us to test for liver function, since some human patients receiving Ceritinib have elevated levels of the liver enzymes alanine aminotransferase (ALT) and aspartate aminotransferase (AST) (Cooper et al., 2015, Duarte et al., 2021). While we failed to detect significant differences in AST concentrations across treatment groups (Fig. 4L), there was a modest but significant increase (p<0.05) in the levels of ALT in mice treated with Cer but not Com (Fig. 4M), mirroring results with GDF-15 (Fig. 4K). Together, these results suggest that a low level of liver toxicity accompanies treatment with Ceritinib, but since the effect is modest and is abolished by co-administration with Semaglutide it cannot explain the dramatic reduction in body weight that is observed (Fig. 4D).

### Ceritinib reduces food intake, but has no effect on energy expenditure

Having ruled out toxicity as the major driver of weight loss, we wondered what mechanisms might be responsible. Metabolic phenotyping studies in mice deficient in *Alk* throughout development suggested that their resistance to DIO was due to increased energy expenditure but not reduced food intake (Orthofer et al. 2020). To test if acute inhibition of Alk by Ceritinib would have a similar effect, we individually housed DIO mice in Promethion metabolic cages to monitor continuous food intake, energy expenditure, and body weight. In these cages, we injected mice daily with Veh, Sem, Cer, or Com for 5 days (Fig. 4N), and observed a similar trajectory of body weight loss (Fig. 4O) as seen previously (Fig. 4D). Relative to the body weights of Veh-treated mice at D5, we observed a significant weight reduction in mice treated with Sem (11.90 ± 0.69%; p<0.001), Cer (13.70 ± 1.86%; p<0.001), and Com (13.53 ± 1.38%; p<0001).

Unexpectedly, indirect calorimetry revealed no significant differences in energy expenditure between any of the treatment groups, regardless of whether we tested for these effects by ANCOVA (Fig. 4P) or normalisation to body weight (Fig. 4Q). However a similar analysis revealed a significant decrease in cumulative food intake in mice treated with Sem (p<0.01), Cer (p<0.001), or Com (p<0.001) (Fig. 4R,S). These effects were dramatic: the food intake of mice treated with Cer or Com was only about a third of Veh controls. These results were consistent across two independent cohorts, and suggest that Ceritinib treatment induces weight loss by suppressing food intake rather than by affecting energy expenditure.

### Ceritinib’s effects on gene expression in white adipose tissue

Since Ceritinib treatment was associated with significantly reduced fat mass, we asked what transcriptional changes are seen in white adipose tissue (WAT), which was reported to have an upregulated lipolysis program in Alk-deficient mice (Orthofer et al. 2020). We therefore examined epididymal WAT (eWAT) from mice treated for 5 days with Veh, Sem, Cer, or Com by RT-qPCR and observed that Cer (p<0.001) and Com (p<0.01) significantly reduced the mRNA expression of peroxisome proliferator-activated receptor γ (*Pparg*), a master regulator of adipogenesis and lipid storage (Fig. 5A). Treatment with Cer also significantly downregulated the mRNAs of CCAAT enhancer binding protein α (*Cebpa*) and perilipin 1 (*Plin1*). These genes play key roles in adipocyte maturation, triglycerides synthesis, and lipid storage (Z. Sun et al. 2013; Zhang et al. 2018). Furthermore, following 5 days of drug treatment, adiponectin (*Adipoq*) was downregulated in the Sem (p<0.05), Cer (p<0.001), and Com (p<0.01) groups, possibly as a consequence of *Pparg* and *Cebpa* downregulation (Qiao et al. 2011). Consistent with the reduction of circulating leptin levels (Fig. 4H), we also found a significant downregulation of *Lep* mRNA in mice treated with Sem (p<0.05), Cer (p<0.001), and Com (p<0.001). Alk-deficient mice fed either a control or HFD were reported to upregulate genes involved in fatty acid oxidation including *Pgc1α*, *Pdk4*, and *Acox1* in eWAT (Orthofer et al. 2020). In mice treated with Cer for 5 days, we found a significant upregulation of *Pdk4* (p<0.01) mRNA expression, but *Pgc1α* and *Acox1* were not significantly changed in eWAT. Together, these results are consistent with the actively reducing fat mass observed in Ceritinib-treated mice, but these are unlikely to drive the observed reduction in food intake.

**Figure 5.**
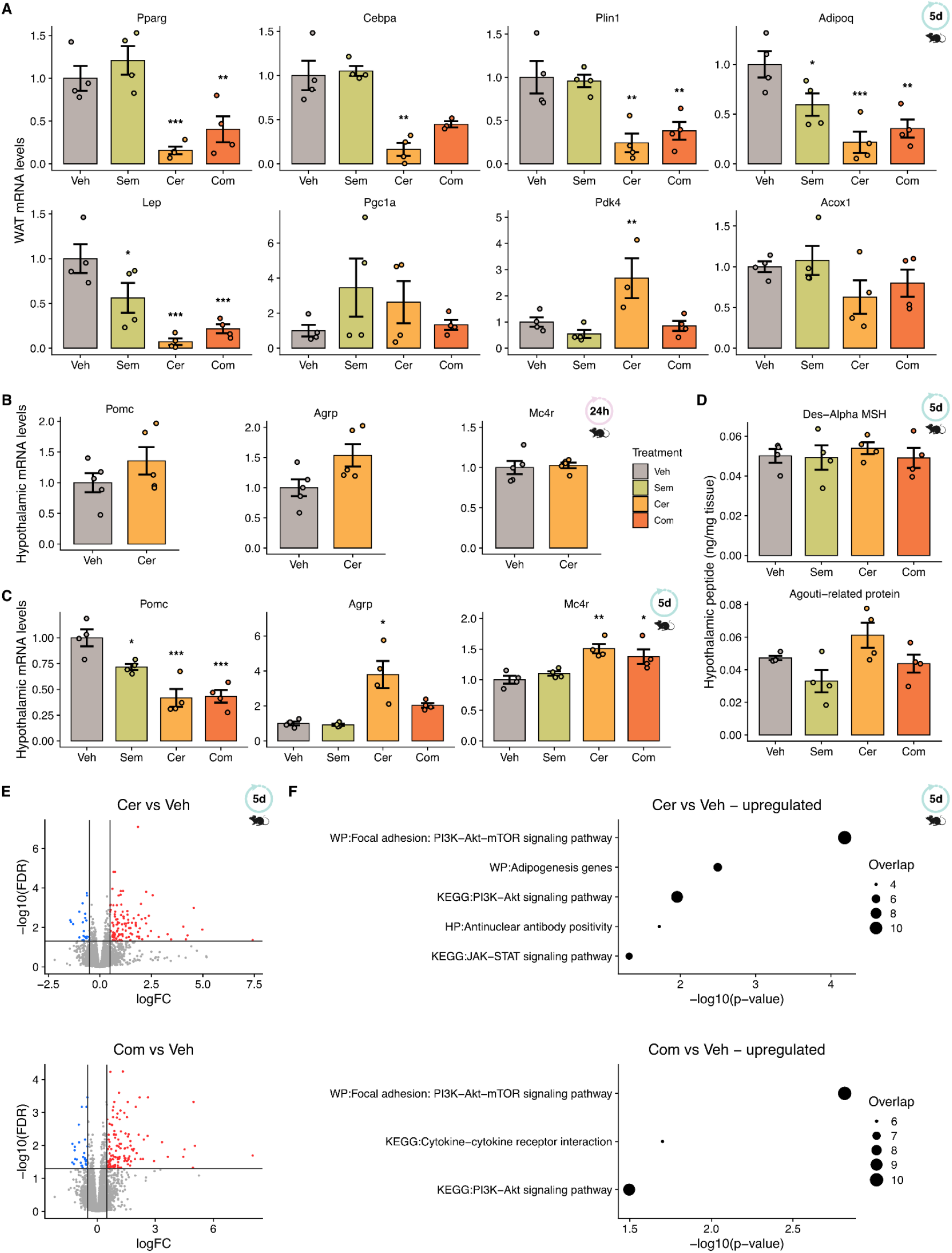
Ceritinib treatment upregulates hypothalamic PI3K/AKT signalling and *Glp1r* expression. **A)** Effects of vehicle (Veh), Semaglutide (Sem), Ceritinib (Cer), and Ceritinib + Semaglutide (Com) on gene expression in epididymal white adipose tissue (n=4/group). **B,C)** 24 hours after Cer or Veh treatment (B), genes in the melanocortin pathways are not significantly affected (n=5/group), but after 5 days days of treatment (C), changes in gene expression are consistent with homeostatic responses to negative energy balance rather than a causal involvement with decreased food intake (n=4/group). **D)** Targeted quantitative peptidomic analysis of the mouse hypothalamus did not reveal significant changes likely to drive reduced food intake (n=4/group). **E, F)** After 5 days of gene expression, Cer or Com treatment induces significant changes in hypothalamic gene expression relative to Veh (E) that consistently involve upregulation of the PI3K/AKT pathway (F, n=4/group). Data represent mean ± SEM; *p<0.05, **p<0.01, ***p<0.001.

### Ceritinib does not appear to potentiate melanocortin signalling

We reasoned that ALK likely acts on the hypothalamus to regulate food intake since it was enriched in hPOMC neurons and earlier studies implicated some of its action in the hypothalamus (Orthofer et al. 2020; Ahmed et al. 2022). Specifically, we hypothesise that the observed reduction in food intake might be due to melanocortin signalling, which could be achieved by the increased expression of *Pomc* or its receptor *Mc4r*, or the decreased expression of the endogenous MC4R antagonist *Agrp*, at the gene or peptide levels. We therefore first tested the effects of 24 hour treatment with Veh or Cer on hypothalamic gene expression in a cohort of DIO mice and observed no significant changes in any of these genes, although there was a trend toward increased *Pomc* expression (Fig. 5B). Next, we examined the effects of drug treatment for 5 days on gene expression in the hypothalamus of DIO mice. We observed a significant downregulation of *Pomc* mRNA in the Sem (p<0.05), Cer (p<0.001), and Com (p<0.001) groups while conversely a significant upregulation of *Agrp* (p<0.05) in the Cer group (Fig. 5C). These results may suggest a homeostatic response to negative energy balance. We also observed an upregulation of *Mc4r* in the Cer (p<0.01) and Com (p<0.05) groups (Fig. 5C). Since gene expression may not reflect relevant changes seen at the protein level, we performed targeted quantitative peptidomics for candidate neuropeptides involved in hypothalamic appetite regulation, but did not observe significant differences in the POMC-derived peptides α-MSH or in Agrp (Fig. 5D). Overall, these results do not support our hypothesis that Alk inhibition increases melanocortin tone and suggest a different mechanism of action.

### Ceritinib upregulates hypothalamic Pi3k/Akt signalling

To identify Ceritinib’s potential mechanisms of action, we performed bulk RNA sequencing of the hypothalamus from mice treated for 5 days with Veh, Sem, Cer, or Com. We identified 246 significantly upregulated and 132 downregulated genes (absolute log2(fold-change) > 0.5 and FDR < 0.05) in the hypothalamus in Cer vs Veh, and similarly found 308 significantly upregulated and 159 downregulated genes when comparing Com and Veh (Fig. 5E). Many of these differentially expressed genes were shared between Cer and Com groups, suggesting that Ceritinb treatment drove most of the observed gene expression changes (Table S3A,B). Gene ontology (GO) analysis did not suggest enrichment in pathways indicative of non-specific toxicity or stress, but instead revealed that the Pi3k/Akt/Mtor signalling pathway was significantly upregulated (analytical adjusted p-value < 0.05) in both the Cer and Com groups (Fig. 5F) and that cholesterol biosynthesis and metabolism were significantly downregulated in the Cer group (Table S3C,D). Since Alk signals via the Pi3k/Akt pathway and Ceritinib inhibits Alk signalling, these results seem paradoxical. However, since Alk shares signal transduction pathways including Pi3k/Akt with several other metabolically important receptors expressed in the same cell types, such as the insulin receptor (Insr), leptin receptor (Lepr), Mc4r, or Glp1r, we hypothesised that inhibiting Alk signalling might potentiate signalling via these receptors.

### Ceritinib potentiates GLP1R signalling in vivo and in vitro

To test this hypothesis, we first asked whether administration of Ceritinib for 24 hours could alter the expression of candidate receptors *in vivo*. In the mouse hypothalamus, the expression of *Insr* and *Lepr* mRNAs were unchanged, but we observed a significant increase (p<0.01) in *Glp1r* (Fig. 6A). This significant increase in hypothalamic *Glp1r* expression was maintained in mice treated for 5 days with Cer (p<0.01) (Fig. 6B). Additionally, we also observed an increase in *Lepr* mRNA in the Cer and Com (p<0.01) groups while no change was detected for *Insr*. To test whether these effects could be replicated in human hypothalamic cells, we treated them with Ceritinib for 24 hours (Fig. 6C) and found a significant upregulation of *INSR* (p<0.001), *LEPR* (p<0.01) and in particular *GLP1R (p<0.01)* whose mRNA expression levels were nearly tripled (Fig. 6D). We therefore hypothesised that the increased expression of *GLP1R* upon Ceritinib treatment might render cultures more sensitive to subsequent administration of Semaglutide. To investigate this, we incubated cultures of hypothalamic neurons with Cer or Veh for 24 hours, and used calcium imaging to test their responses to 200 nM Semaglutide (Fig. 6C). We found that changes in intracellular calcium concentrations (ΔF/F) in response to Sem were significantly (p<0.05) potentiated by pre-treatment with Cer, as determined by taking the mean area under the curve (AUC) after Sem administration (Fig. 6E,F). These results were consistent across three replicate experiments, and suggest that the Ceritinib-induced increase in *GLP1R* expression observed across both mice and human cells potentiates responses to endogenous GLP-1 and contributes to its suppression of food intake in DIO mice and the additive effects observed with combined drug treatment.

**Figure 6.**
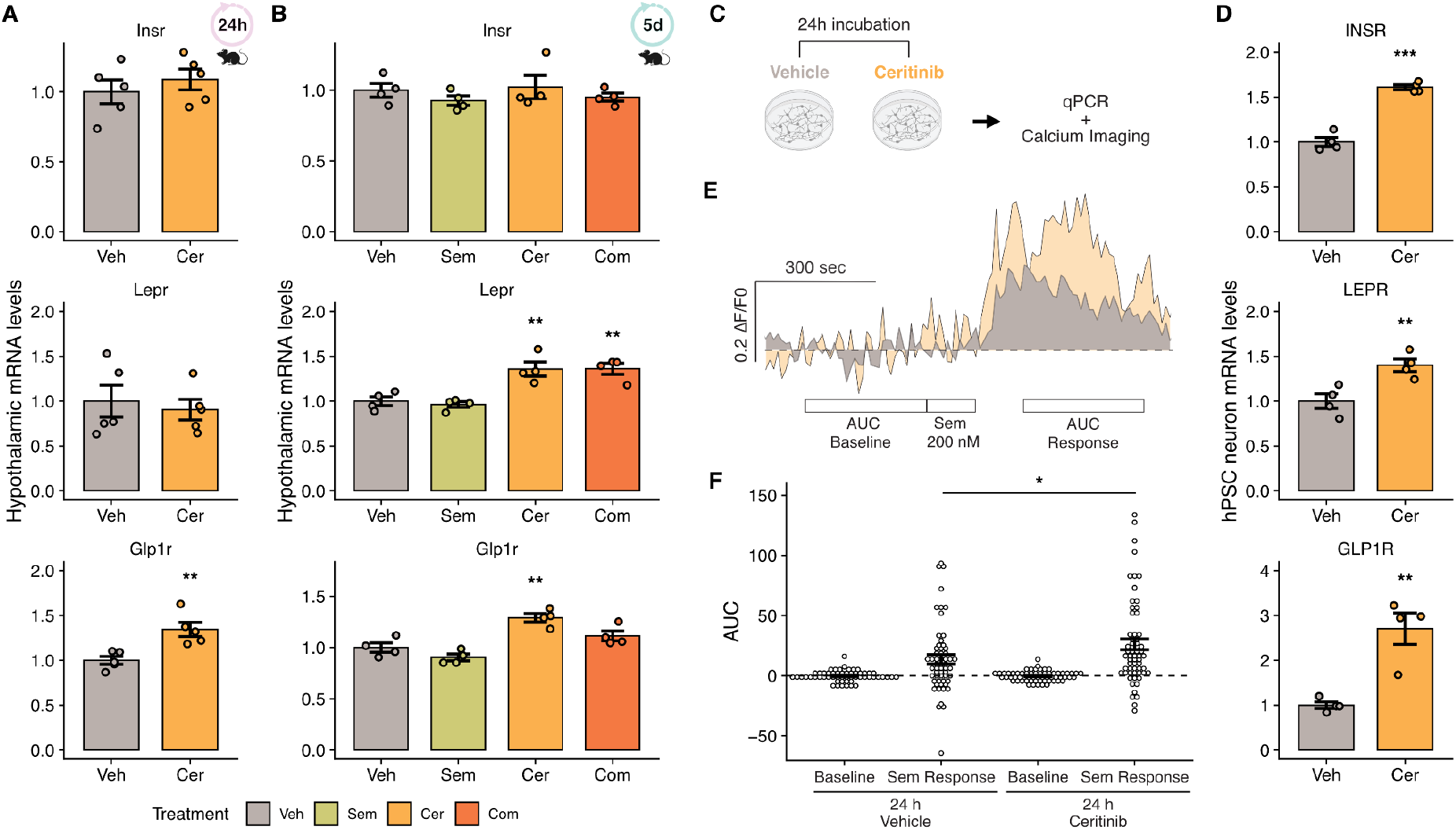
Ceritinib potentiates GLP1R signalling. **A, B)** After both 24 hours of treatment with Veh or Cer (A) or 5 days of treatment with Veh, Sem, Cer, or Com, *Glp1r* is consistently upregulated in Cer-treated animals. **C)** Experimental schematic of 24 hour incubation with Veh or Cer in human hypothalamic cultures, followed by qPCR, or by administration of Semaglutide (Sem) and calcium imaging. **D)** Cer significantly increases the expression of several genes of interest, notably *GLP1R* (n=4/group). **E)** Representative calcium imaging traces of neurons incubated with Vehicle (grey) or Ceritinib (orange), and treated with Sem. **F)** Responses to Sem are significantly (p<0.05) potentiated by pre-treatment with Cer (n=62, N=3) relative to vehicle (n=57, N=3). Data represent mean ± SEM (qPCR) and mean with 95% CI (calcium imaging); *p<0.05, **p<0.01, ***p<0.001.

## Discussion

Given the central role of POMC neurons in regulating food intake, we hypothesised that targeting genes enriched in this cell population would regulate their activity and might reveal new pharmacological strategies for treating obesity. We tested this hypothesis in hPSC-derived POMC neurons and identified 13 candidate genes targeted by drugs approved for use in humans, including ALK. We found that treatment with the potent and selective ALK inhibitor Ceritinib dramatically reduced food intake and body weight in mice. Here, we discuss the insights gleaned from our experimental model system and the clinical implications of our findings.

Using scRNAseq of hypothalamic cultures generated from five genetically distinct hPSC cell lines and grown under four different media conditions, we showed robust differentiation of these to predominantly anterior ventral hypothalamic cell types that transcriptionally resembled primary neonatal hypothalamic cells, including POMC neurons. We also observed the production of ependymal-like cells and tanycyte-like cells, including cell clusters enriched in markers indicative of beta-tanycytes that line the ventral walls of the third ventricle and are thought to play important roles in the trafficking of leptin and other factors across the BBB (Balland et al. 2014). Future studies of these hPSC-derived cells may shed light on these transport mechanisms and reveal how tanycytes functionally interact with other hypothalamic neuron populations. Overall, the transcriptional similarity of hPSC-derived and neonatal human POMC neurons suggested that our model system might reveal insights into obesity mechanisms when integrated with human genetic data.

We found that hPOMC neurons were enriched in GLP1R and robustly responded to the GLP1R agonist Semaglutide. We also explored ALK signalling in hPOMC neurons and found that acute administration of its agonist ALKAL2 leads to a rapid increase in intracellular calcium concentration that is likely mediated at least in part by PI3K signalling, whereas longer-term administration with the ALK inhibitor Ceritinib upregulated the PI3K/AKT pathway. We propose that these longer-term effects are due at least in part to the transcriptional upregulation of genes such as GLP1R that we consistently observed both *in vitro* and *in vivo*, and whose protein products signal via the PI3K pathway.

We found that Ceritinib-induced weight loss was associated with decreased food intake but unchanged energy expenditure, whereas mice constitutively lacking Alk were reported to have unaltered food intake and increased energy expenditure (Orthofer et al. 2020). Since Alk plays roles in both brain development and adult function, we propose that this discrepancy may be explained by the altered development of Alk^-/-^ mice and/or the activation of compensatory pathways that are not engaged upon the acute inhibition of Alk in the adult brain. It is unlikely that the effects of Ceritinib administration can be explained by non-specific toxicity or nausea for several reasons. First, these factors should cause reduced food intake and body weight in both lean and DIO mice, but lean mice treated with Ceritinib gained weight at the same rate as vehicle-treated controls. Second, we did not observe elevations in the stress marker CRP, or observe transcriptional signatures of stress in the brains of mice in any treatment group, which instead suggested alterations in the signalling pathways engaged by Alk. Third, although plasma concentrations of GDF15 and ALT were moderately but significantly elevated in mice treated with Ceritinib alone, co-administration of Semaglutide completely normalised them to levels seen in vehicle-treated mice despite an even greater reduction in mouse body weight. We speculate that the ability of semaglutide to reduce markers of liver toxicity may prove beneficial to cancer patients taking Ceritinib.

In our view, it is premature to suggest that Ceritinib or other ALK inhibitors could be a safe and effective treatment for metabolic disease in humans, but if animal studies suggest that the drug side effects can be mitigated, then the long-term inhibition of Alk signalling could be benign. Alk knockout mice are fertile, have no obvious developmental or cognitive defects, and show elevated hippocampal neurogenesis and improved performance in stress and memory tests (Bilsland et al. 2008). Furthermore, female fruit flies lacking Alk signalling live longer with preserved locomotor function and are relatively resistant to stress (Woodling et al. 2020). Overall, our studies have shown that hPSC-derived hypothalamic neurons are a powerful model system to connect human genetics of obesity to cellular phenotypes and accelerate the development of new therapies for obesity.

## Limitations

hPSC-derived cell types are relatively immature and their transcriptional profiles most closely resemble those observed in human foetal tissue (Nicholas et al. 2013), limiting our ability to study disease-relevant transcriptional signatures and functional properties relevant to the fully mature cells. Despite this, the transcriptional similarity of hPOMC to neonatal pPOMC neurons and their functional responsiveness to GLP-1 and ALKAL2 suggest that they are sufficiently mature to facilitate biological discovery. Furthermore, we anticipate that by improving methods to generate hypothalamic cells (e.g. by transcription factor-based forward programming) and maturing them (e.g. in assembloids) will further enhance their utility as a drug discovery platform and enable the approach we report here to be extended to other hypothalamic cell types of interest.

In this study, we considered genes likely to be associated with obesity by a pipeline that considers GWAS, eQTL, and enhancer mapping data (Kentistou et al. 2023). The causality of genes prioritised in this manner is uncertain, and in the future it would be interesting to test the effects of other genes associated with obesity by population exome and genome sequencing data, as well as genes nominated by familial or cohort studies of severe obesity. Furthermore, for practical considerations, we considered only genes that already have drugs approved for human use, and further prioritised those with published evidence that they penetrate the BBB. The approach could be expanded to include a broader array of small molecule drugs and biologics targeting genes of interest.

Although Ceritinib is a potent and selective ALK inhibitor and the doses we gave to mice (5-10 mg/kg/day) are similar to those used in humans (∼6 mg/kg/day), we cannot formally exclude the possibility that some of the effects we observed *in vitro* and *in vivo* might be attributable to off-target effects at the effective concentrations that reach the brain and other organs. We demonstrated that Ceritinib treatment increased hypothalamic PI3K/AKT signalling, increased the expression of *GLP1R* in human hypothalamic cultures and *Glp1r* in the mouse brain, and enhanced the responses of hypothalamic neurons to Semaglutide. However, ALK is widely expressed in the nervous system and is also expressed at lower levels in other tissues, so other brain regions could contribute to the observed Ceritinib-induced appetite suppression. In the future, it will be interesting to explore its effects in other candidate brain regions such as the nucleus tractus solitarius (NTS) of the hindbrain.

While our results with Ceritinib demonstrate significant weight loss that could in theory be translated rapidly into human trials, there are several reasons for caution. Since pharmacological management of body weight requires long-term treatment, the benefits of increased metabolic health must clearly outweigh the potential side effects of the drug, especially when given for longer periods. At the doses we used, side effects such as liver toxicity are observed in some cancer patients, suggesting that further optimisation of drug dosage or exploration of other ALK inhibitors is necessary.

## Materials & Methods

### Human pluripotent stem cell (hPSC) lines

All human embryonic stem cell (hESC) and human induced pluripotent stem cell (hiPSC) lines used in this study were previously generated and reported to be derived from material obtained under informed consent and appropriate ethical approvals. Cell lines used in this study included KOLF2.1J (Passage 7-16), NCRM5 (Passage 16-21), NN0004297 (Passage 29), NN0003932 (Passage 44), PGP1 (Passage 28), and a POMC-GFP reporter line (as described below) generated on the HUES9 background (Passage 12-18 after reporter generation). Gene editing and other work with these lines as part of this study was performed under approved protocols from our respective institutions.

### Generation of a POMC-GFP knock-in reporter cell line

The POMC-GFP reporter cell line was generated using CRISPR/Cas9 genome editing of the human embryonic stem cell line HUES9 as previously described (Santos et al. 2016). Briefly, HUES9 hESCs were dissociated to a single cell suspension and nucleofected with recombinant spCas9 protein, a CRISPR sgRNA sequence targeting the STOP codon of the endogenous human POMC gene and cutting right before this codon (CCTACAAGAAGGGCGAGTGA). The targeting vector contained a 5’ homology arm of endogenous coding human POMC sequence that terminated just upstream of the TGA stop codon constituting the final three bases of the CRISPR and instead replaced it in-frame with the T2A ribosomal skipping sequence and a green fluorescent protein (GFP) sequence with its own stop codon and polyadenylation (polyA) site, followed by a floxed RFP-T2A-puromycin-polyA drug selection cassette under control of a constitutive promoter. After nucleofection and recovery, cells were selected with puromycin and individual colonies were picked, expanded, and screened by PCR for the fidelity of their integration in the POMC locus. We selected clone 64 that gave a clear PCR product spanning the endogenous POMC coding sequence into the genomics sequence downstream of the 3’ homology arm, indicating correct integration, and electroporated this clone with a Cre-expressing construct to remove the floxed puromycin selection cassette. Individual colonies were then picked, expanded, and analysed by PCR to confirm both the removal of this cassette and the absence of PCR products from the targeting vector backbone indicative of additional integrations elsewhere in the genome. Clones 6E, 7A, 7B, 7E, and 10H gave PCR products indicative of correct targeting only at the endogenous POMC locus and were differentiated into hypothalamic neurons. Upon differentiation, clone 10H consistently yielded >10% brightly green fluorescent cells immunopositive for POMC. Microarray analysis of this clone confirmed the absence of culture-acquired copy number variants, and it was therefore selected for further use in this study.

### HPSC maintenance

hPSC lines were maintained under feeder-free conditions without antibiotics in mTeSR1 (STEMCELL technologies) or StemFlex media (Thermo Fisher Scientific) in humidified incubators at 37 °C and 5% CO2. hPSC lines were seeded on plates coated with Geltrex^TM^ (Thermo Fisher Scientific) at a 1:100 dilution prepared in DMEM/F12 (no phenol red; Thermo Fisher Scientific). mTeSR1 or StemFlex mediums were kept at 4°C for no longer than 1 month after supplement addition, and only required volumes of media were warmed to 37 °C immediately before use and culture medium was replaced every 24-48 hours. For all cell lines, colonies had clearly defined borders and lacked differentiated cells when viewed under a phase contrast microscope. Cultures were passaged when they reached 60-80% confluence by a brief wash with DPBS without calcium and magnesium (Thermo Fisher Scientific) followed by an approximately 5 minute incubation with EDTA (0.5 mM, pH 8.0, Thermo Fisher Scientific) for maintenance or TrypLE (Thermo Fisher Scientific) for differentiation. A single-cell suspension was then achieved by gentle mechanical dissociation in media containing 10 µM Rock inhibitor Y-27632 (DNSK International). Cell suspensions were then pelleted by centrifugation at 160 *g* for 3 minutes, resuspended, and replated for maintenance or differentiation in media containing 10 µM Rock inhibitor as described below.

### Directed differentiation to hypothalamic progenitors

hPSCs were differentiated to hypothalamic neurons by directed differentiation as previously described (Merkle et al. 2015; Kirwan, Jura, and Merkle 2017; H.-J. C. Chen et al. 2023). Briefly, hPSCs were dissociated to a single-cell suspension with TrypLE, pelleted and resuspended in media containing 10 µM Rock inhibitor, followed by cell counting on a Countess II automated cell counter. Cells were then plated on geltrex (1:100) coated 12-well plates at a density of 1 x 10^5^ cells / cm^2^ in the same medium. The following day, cultures were washed once with DPBS (without calcium and magnesium) and switched to N2B27 media, consisting of 500 mL Neurobasal-A (Thermo Fisher Scientific), 500 mL DMEM/F12 with GlutaMAX (Thermo Fisher Scientific), 10mL Glutamax (Thermo Fisher Scientific), 10 mL sodium bicarbonate (Thermo Fisher Scientific), 5 mL MEM Non-essential Amino Acids (Thermo Fisher Scientific), 1 mL 200 mM ascorbic acid (Sigma-Aldrich) 10 mL 100x penicillin-streptomycin (Thermo Fisher Scientific), 20 mL 50x B27 supplement (Thermo Fisher Scientific), 10 mL 100x N2 supplement (Thermo Fisher Scientific). Small molecule modulators of signalling pathways were used to direct hypothalamic differentiation as previously described (H.-J. C. Chen et al. 2023). Specifically, initial concentrations of 100 nM LDN-193189 (DNSK International), 10 mM SB431542 (DNSK International), and 2 mM XAV939 (DNSK International) were gradually reduced from day 0 (D0) to D10 of differentiation, and SAG (Fisher Scientific) and purmorphamine (Fisher Scientific) were each added to a final concentration of 1 mM from D2-D8.

### Expansion and maintenance of primary astrocyte cultures

Mouse primary astrocyte was purchased from ScienCell Research Laboratories (Carlsbad, CA) and cultured in astrocyte media composed of DMEM/F12 with GlutaMax (Thermo Fisher Scientific), 10% (vol/vol) foetal bovine serum (FBS; Thermo Fisher Scientific), and 2% (vol/vol) penicillin-streptomycin (Thermo Fisher Scientific) at 37 °C in a humidified atmosphere of 5% CO2. Primary mouse astrocytes (passage 0 - 5) were maintained in Geltrex^TM^-coated T75 culture flask (Thermo Fisher Scientific) and re-plated at 5.7 x 10^4^ cells/cm^2^ to 1 x 10^5^ cells/cm^2^ 3-5 days prior to seeding hypothalamic progenitors for the co-culture experiments including immunocytochemistry, whole-cell patch clamp electrophysiology, and single cell RNA sequencing.

### Hypothalamic neurogenesis and maturation

After 14 days of hypothalamic differentiation, hypothalamic progenitors were dissociated with a mixture of TrypLE, papain (Worthington Biochemical), and DNAse I (Worthington Biochemical) with a subsequent 2 washes in N2B27 medium supplemented with 10 µM Rock inhibitor. Following the final centrifugation at 160 *g* for 3 minutes, the cell pellet was resuspended, counted, and replated at a concentration of 1 x 10^5^ cells/cm^2^ in two of the four tested conditions: **(1)** N2B27 medium (see above) containing 10 *µ*g/ml BDNF (Qkine) (NB) and **(2)** co-cultures of hPSC-derived neurons with primary mouse astrocytes in N2B27 medium (NBA). For the other two tested conditions: **(3)** Synaptojuice medium 1 (SJ1) followed by Symaptojuice medium 2 (SJ2), both containing 10 *µ*g/ml BDNF (SJ) and **(4)** co-cultures of hPSC-derived neurons with primary mouse astrocytes in SJ medium (SJA), cells were seeded at a concentration of 3 x 10^5^ cells/cm^2^. More specifically, SJ medium was adapted from Kemp and colleagues (Kemp et al. 2016) and hypothalamic cultures were maintained in SJ1 (N2B27, 2 μM PD0332991 isethionate (Sigma Aldrich), 5 μM DAPT (DNSK International), 370 μM CaCl_2_ (Sigma Aldrich), 1 μM LM22A4 (Tocris), 2 μM CHIR99021 (Cambridge Bioscience), 300 μM GABA (Tocris), and 10 μM NKH447 (Sigma Aldrich) for 1 week. Cultures were then switched to SJ2 (N2B27, 2 μM PD0332991 isethionate, 370 μM CaCl_2_, 1 μM LM22A4, and 2 μM CHIR99021) for the rest of the maturation period until at least 40 days post-differentiation as per described previously (H.-J. C. Chen et al. 2023). Cultures were fed every 2 days by a complete media change, and all media were supplemented with 1 µg/mL of laminin (Sigma) to help maintain cell attachment.

### Immunocytochemistry and imaging

Hypothalamic progenitors were re-plated onto PhenoPlate^TM^ 96-well microplates (Perkin Elmer) and hypothalamic cultures at 33-40 days post-differentiation were fixed in 4% paraformaldehyde (Thermo Fisher Scientific) for 10 minutes, rinsed three times with Tris-buffered saline containing 0.1% Triton X-100 (TBS-T; Sigma-Aldrich). Cells were then incubated with primary antibodies including anti-MAP2 (Abcam; 1:2,000) and anti-αMSH (A1H5, detects epitopes in the αMSH region of POMC and developed by Professor Ann White; 1:5000) diluted in TBS-T containing 1% normal donkey serum (NDS; Stratech) overnight at 4 °C. Cells were washed three times with TBS-T and incubated with secondary antibodies combining donkey anti-mouse AF555 (Thermo Fisher Scientific) and donkey anti-chicken AF647 (Thermo Fisher Scientific) diluted in TBS-T with 1% NDS for 2 hours at room temperature on an orbital shaker. Cells were washed three more times with TBS-T, and 300 µM of 4′,6-diamidino-2-phenylindole (DAPI; Thermo Fisher Scientific) was used to stain cell nuclei. Image acquisition was performed using the Perkin Elmer Opera Phenix Plus High-Content Screening System with a 20x water objective. A total of 21 fields of view per well with 5 z-stacks per field at 0.8 µm intervals were imaged.

### Imaging analysis using Harmony

Image analysis of hypothalamic cultures stained with DAPI (DAPI 405 channel), αMSH (POMC; Alexa 555 channel), and MAP2 (Alexa 647 channel) was performed using the Harmony High-Content Imaging and Analysis software (Perkin Elmer, v4.9). Specifically, an automated analysis pipeline was developed to first detect DAPI positive nuclei and quantify the nuclear area and number per field. Next, a fixed-intensity threshold was applied to every image in the 647/MAP2 channel to exclude background noise, followed by 647 positive MAP2 object detection as primary objects and the measuring of neuronal number and area. The 555 channel was analysed using the same approach for POMC-positive staining. A mask was then created using the 647/MAP2 channel and applied to the 555/POMC channel to calculate the percentage of POMC-positive neurons out of the total number of neuronal MAP2 (see supplementary material for detailed analysis sequence).

### Whole-cell patch clamp electrophysiology

Hypothalamic progenitors differentiated from the HUES9-POMC-GFP cell line were re-plated onto 24 mm round glass coverslip (VWR) that has been pre-treated with 5 M hydrochloric acid and double coated with both 0.01% poly-L-ornithine hydrobromide (Sigma-Aldrich) and Geltrex^TM^. Whole-cell patch-clamp recordings were performed as described previously (Bilican et al. 2014). Green fluorescent neurons derived from the HUES9-POMC-GFP cell line maintained in NB, SJ, or SJA, for 40+ days post-differentiation were selected for patching. Recordings were made in the presence of blockers of synaptic transmission (PTX, 50 µM; strychnine, 20 µM; APV, 100 µM; CNQX, 30 µM) to enable characterisation of intrinsic electrical properties without the influence of other cells in the culture.

### Cell dissociation for single cell RNA sequencing

Hypothalamic progenitors were re-plated onto 6-well or 12-well polystyrene plates (Corning) for scRNAseq experiment. Hypothalamic cultures at 40 ± 1 days post-differentiation were gently washed twice with calcium and magnesium free DPBS (Thermo Fisher Scientific) before adding dissociation solution consisting of TrypLE™ Express (Thermo Fisher Scientific) supplemented with papain (Worthington) and 30 µM actinomycin D (Sigma) to block transcription as previously described (Chen & Merkle, 2020; protocol io). Cells were incubated at 37 °C for up to 5 minutes before aspirating the dissociation solution followed by addition of wash medium (N2B27) supplemented with 10 µM ROCK inhibitor, 33 μg /mL DNase I (Worthington), and 30 µM actinomycin D. The cells were dissociated using a P1000 pipette and subsequently passed through a 40-µm Flowmi® cell strainer (Merck) into DNA Lo-bind Eppendorf tubes (Eppendorf). After centrifugation at 200 *g* for 3 minutes and another 2 washes, the cells were resuspended in DPBS (no calcium and magnesium) containing 0.04% BSA (Sigma Aldrich) and 30 µM actinomycin D. Cells were passed through the 40-µm Flowmi® cell strainer again, stained with 0.4% Trypan Blue, counted using the Countess^TM^ cell counting chamber slides (Invitrogen), and concentrations were adjusted to 4.26 x 10^5^ cells per mL. An aliquot of 47 µL for each cell line and condition to achieve a targeted cell count of 20,000 was submitted to Cancer Research UK (CRUK) Genomic Core for cDNA library preparation.

### Single-cell library generation and sequencing

Libraries were prepared using the 10x Genomics Chromium Controller in conjunction with the single-cell 3′ v3 & 3.1 kits (10x Genomics). The cDNA synthesis, barcoding, and library preparation were carried out according to the manufacturer’s instructions. Libraries were sequenced in CRUK on a Novaseq 6000 sequencer (Illumina).

### scRNA-seq data processing

Raw sequencing libraries were processed using STARsolo (version 2.7.9) (Kaminow, Yunusov, and Dobin 2021) using recommended parameters for producing output matching 10x Genomics’ Cell Ranger platform. Reads were aligned and quantified to a ‘barnyard’ reference genome composed of human (GRCh38, Ensembl release 98 with gencode annotation release v32), mouse (GRCm38, Ensembl release 98 with gencode annotation release vM23) and GFP-reporter references, built using the scripts provided by 10x Genomics and 10x Genomics Cell Ranger mkref function (version 6.0.1). Droplets containing captured cells were called using emptyDrops function from the DropletUtils R package (Lun et al. 2019), with UMI threshold calculated adaptively per sample based on the mean and standard deviation of the background counts distribution; and an FDR of 0.001.

### Cell species calling and cell line demultiplexing

Cell species identity for all samples was determined following a modified cell species assignment method from 10x Genomics. Briefly, cells were identified as either human or mouse cells based on the proportion of unique molecular identifier (UMI) content belonging to human and mouse genes, with cells containing >90% UMI from a single species being identified as belonging to that species. Cells containing <=90% UMI from a single species or cells with more than the batch-specific threshold of UMI counts from the other species were excluded from downstream analysis. Cells identified as human were stripped of UMI counts belonging to mouse for downstream analysis. In conditions where genetically distinct hPSCs were multiplexed per condition prior to 10x library generation, cell line identity was inferred based on the genotype using 10x Genomics Vartrix (v1.1.22) and vireo SNP tools (v0.5.6) (Y. Huang, McCarthy, and Stegle 2019). Variant-count matrices were generated with Vartrix using aligned reads from STARsolo and variant call format (VCF) file with information on each sub-line genotype (Pantazis et al. 2022) as inputs. Cell line identity was inferred using the Vireo tool with the variant-count matrices and variant information. Since the NN0003932 and NN0004297 cell lines were derived from the same donor, we considered reads from either of these as a single genetically distinct cell line, denoted as NN_combined.

### Quality control of single cell sequencing data

Cell filtering was performed for each sample to remove low quality and outlier cells based on quality control metrics of total UMI content, number of detected features/genes, fraction of mitochondrial and ribosomal content, as well as cell line assignment. Comparisons of quality control metrics across samples identified one sample (first SJ differentiation batch) with double the UMI content distribution compared to other samples due to the lower cell capture efficiency and subsequently deeper sequencing depth of the sample. Downsampling of UMI counts was performed on the deeply sequenced sample using the downsampleMatrix function from DropletUtils R package to match the sequencing depth of the next highest sample. Cells were then discarded if their UMI content or number of features were more than 3 median absolute deviation (MAD) away from the median, or if the fraction of mitochondrial or ribosomal content of the cell was higher than 4 MAD from median.

Cell gene expression was normalised using logNormCounts function from the scran R package (Lun, McCarthy, and Marioni 2016), with size normalisation factors computed using the computeSumFactors function. Clustering information for computing size factors were calculated using the quickCluster function, which utilises the walktrap algorithm for cluster assignment (Pons and Latapy 2005). The resulting normalised gene expression levels were then evaluated by comparing the distribution of total normalised expression of stable expressed genes per cell across samples. For more details, please see the documentation in the open-access code that accompanies this manuscript.

### Doublet detection

Doublet detection was performed for each sample individually as well as jointly across all samples. For sample level doublet detection, cells were initially identified as ‘hybrid doublets’ in each sample using the cxds_bcds_hybrid function from scds R package (Bais and Kostka 2019), with estNdbl parameter set to true. This was followed by a cluster-based ‘guilt-by-association’ doublet identification approach, where clustering was first done for each cell (see below) and then repeated for each identified cluster to form smaller, fine-grained clusters. Doublet metrics were then calculated for each sub-cluster based on the fraction of ‘hybrid doublets’ and ‘sub-line doublets’ identified in the cell line demultiplexing step. Cells belonging to ‘guilty’ sub-clusters with either enriched ‘hybrid doublets’ (> 3 MAD away from the median) or ‘sub-line doublets’ (> 6 MAD away from the median) were further identified as doublets. For joint detection of doublets across all samples, samples were first batch corrected (see below) into a single dataset, followed by two rounds of clustering to obtain sub-clusters for identification of cross-sample ‘guilt-by-association’ doublets. As before, cells belonging to ‘guilty’ sub-clusters with enriched per-sample doublets (> 6 MAD away from the median) were then identified as ‘cross-sample doublets’. Finally, cells which were identified as either ‘hybrid doublets’, ‘sub-line doublets’, ‘per-sample doublets’ or ‘cross-sample doublets’ were excluded for further downstream analysis.

### Batch correction and dimensionality reduction

Batch correction was performed across samples to correct for technical variability between samples and to integrate the samples into a single dataset using the fastMNN function from the Batchelor R package (Haghverdi et al. 2018) on the first 50 principal components (PC). Highly variable genes used for computing the principal components were selected using the modelGeneVar function from the scran R package by filtering for all genes with higher variance from the mean-variance curve fitted on the normalised gene expression profile. Both mitochondrial and ribosomal large and small subunit genes were excluded from mean-variance curve fitting due to their high variability and expression. The density.weights option was set to false to avoid overfitting of the mean-variance curve for genes with high variability and expression. Samples were integrated hierarchically based on the experimental batch and differentiation media. Uniform Manifold Approximation and Projection (UMAP) 2D embedding (McInnes, Healy, and Melville 2018) was then calculated from the corrected PC space for dataset visualisation with spread set to 1 and minimum distance set to 0.4.

### Clustering and annotation

Clustering was performed using the Louvain community detection-based method. Shared nearest-neighbour graphs were first constructed from the 50 corrected PC space, followed by clustering with the cluster_louvain function from the igraph R package (Csárdi et al. 2023). Manual annotation was performed for each cluster based on a list of curated genes known to be relevant to hypothalamic differentiation and cell identity, and genes most highly differentially expressed in each cluster.

### Cross-sample quality control and clustering by media type

After joint processing of all samples, the PGP1 cell line was identified to behave differently compared to other cell lines, particularly for cells matured in SJ media. Therefore, cells from this genetic background were removed and the remaining cells were again batch-corrected and clustered as above. Low quality cells were subsequently identified from clusters with low UMI content and number of features, and/or from clusters with high fraction of mitochondrial and ribosomal content and removed. Finally, since the maturation media used strongly influenced the clustering of jointly processed data, final batch correction and clustering was performed on groups of samples cultured in the same media (NB and NBA, or SJ and SJA).

### Comparison of hPSC-derived and primary human neonatal hypothalamic neurons

To assess the transcriptional similarity of hPSC-derived and primary hypothalamic neurons, scRNAseq data from hPSC-derived neurons were integrated with a published dataset of single-nucleus RNAseq data (W.-K. Huang et al. 2021) of human neonatal hypothalamic neurons. The primary human neonatal hypothalamic data was first subset to cells which have been classified as arcuate neurons, and the gene expression was then re-normalised as described above. The two datasets were then subset to genes common between the datasets and normalised using the multiBatchNorm function from the scran R package to account for the different library sizes and single cell capture technology used. Integration of the two datasets was performed with fastMNN as described in the batch correction section above using the published highly variable gene list. Label transfer of primary hypothalamic neurons onto hPSC-derived neurons was performed by finding the 10 nearest primary hypothalamic neurons ‘neighbours’ for each hPSC-derived neurons using the queryKNN function from the BiocNeighbors R package on the integrated space, followed by assigning the most common primary hypothalamic neuron cell type from the neighbouring cells.

### Identification of hPOMC enriched genes

For both NB and SJ base media conditions, differential expression analysis was performed between POMC neurons and other neurons generated in the same differentiations to identify genes enriched in POMC neurons. Non-neuronal cells were first removed, and remaining neurons were classified as POMC or non-POMC based on cluster annotations, as described above. For each cell line, gene counts for POMC and non-POMC cells were then summed to create pseudo-bulk gene counts. In the case of KOLF2.1J pseudobulking was separately performed for the two replicate differentiations. Cell lines with less than 10 cells for either label were excluded from further analysis. Moreover, genes with low expression were removed using the filterByExpr function from the edgeR R package (Y. Chen, Lun, and Smyth 2016). Size factors for the normalisation of pseudo-bulk gene counts were calculated using TMM method, followed by estimation of both negative binomial dispersions and quasi-likelihood dispersion. Differential expression testing was then performed using a quasi-likelihood *F*-test in a paired design to identify genes which are differentially expressed in POMC neurons versus other neurons, with significance threshold set to absolute log2(fold-change) > 0.5 and FDR < 0.05.

### Identification of genes associated with childhood and adult obesity

Genes differentially expressed in POMC neurons, as prioritised above, were integrated with common variant genome-wide association study (GWAS) data on body mass index (BMI, up to n=806,834) from GIANT (Yengo et al. 2018) and comparative size at age 10 (SAC, n=444,345) from UK Biobank (Bycroft et al. 2018). To identify independent GWAS signals and prioritise causal genes at each locus, we used the ‘GWAS to Genes’ pipeline as described previously (Kentistou et al. 2023) and summarised below. Briefly, we filtered the BMI GWAS summary statistics and only retained variants with a minor allele frequency (MAF) >0.1%. Quasi-independent genome-wide significant signals were then selected in 1Mb windows and secondary signals at each locus were further selected via conditional analysis in GCTA (Yang et al. 2011). The primary signals were then supplemented with independent (R^2^<0.05) secondary signals and signals were mapped to proximal genes, within ±500kb of each signal. Independent signals and their closely linked SNPs (R^2^>0.8) were annotated if they were non-synonymous or physically colocalised with known enhancers of the identified genes (Nasser et al. 2021). We also calculated gene-level associations (de Leeuw et al. 2015), by using all common non-synonymous variants within a gene. Colocalization analyses between the GWAS and eQTL (GTEx Consortium 2015; Võsa et al. 2021; Qi et al. 2018) or pQTL (Pietzner et al. 2021) data were also performed using SMR-HEIDI (Zhu et al. 2016) and coloc (Giambartolomei et al. 2014). Finally, genes at the associated loci were also prioritised through the PoPS method (Weeks et al. 2020). Candidate causal genes were prioritised by overlaying all of the above information and scoring the strength of evidence observed.

### Gene prioritisation and target selection

Target genes for further validation experiments were selected from hPSC-POMC neuron enriched gene lists based on several prioritisation criteria. First, the enriched gene lists for each base media type were ranked based on the geometric mean of the fold-change ranking and FDR ranking. Second, these ranked lists were compared with obesity-associated genes as described above to identify genes predicted to contribute to obesity by acting in POMC neurons. Third, candidate drugs that could potentially be repurposed to treat obesity and/or type 2 diabetes were identified based on their action on POMC-enriched and obesity-associated genes, and their approved status for use in humans as represented in the Pharos database (v3.9.0, https://pharos.nih.gov, accessed 11 November 2021). Fourth, remaining genes were manually prioritised for testing *in vitro* and/or *in vivo* based on drug availability, likely penetrance of the blood-brain barrier, support of the gene’s role in body weight regulation in the literature, and enrichment of the gene in POMC neurons in both NB and SJ media.

### Generation of ALKAL2 peptide

The ALKAL2 synthetic gene fragment was prepared by Twist Bioscience (California, USA) using the human ALKAL2 protein sequence (residues 85-152, Uniprot Q6UX46-1) as the template for DNA codon-optimization for expression in *E. coli*. The synthetic gene was cloned into a pExp-DsbC expression vector (Addgene #129243) containing an N-terminal His 8-tag fused to a DsbC solubility enhancing tag, followed by a TEV protease cleavage site.

The expression construct was used to transform competent cells OrigamiB(DE3) (Merck Millipore) and grown overnight at 37 °C on LB-agar plates containing 100 *μ*g ml^−1^ of ampicillin. Large-scale expression was performed by inoculating 1 L of 2YT media using colonies collected from LB-agar plates. Protein expression was induced by 400 *μ*M IPTG at OD_600_ of 0.6 – 0.8 and carried out at 15 °C for 20 h. The next day cells were pelleted by centrifugation at 4,000 *g* for 20 min, re-suspended in TBS buffer (50 mM Tris-HCl pH 8.5, 150 mM NaCl) and stored at −80 °C.

The cell pellet from 1 L of bacterial culture was suspended in 20 mL 50 mM Na_2_HPO_4_ pH 8.5, 20 mM Imidazole, 500 mM NaCl, 5 mM GSH, supplemented with protease inhibitor (cOmplete Mini, EDTA-free, Roche) and lysed by sonication. The lysate was centrifuged at 16,000 *g* for 20 min to pellet the insoluble fraction, filtered and incubated with 5 mL PureCube Ni-NTA agarose (Cube Biotech) for 1 h at 21°C. The resin was washed with 5 CV of 50 mM Na_2_HPO_4_ pH 8.5, 20 mM Imidazole, 500 mM NaCl and eluted in 5 mL fractions with 50 mM Na_2_HPO_4_ pH 8.5, 500 mM Imidazole, 500 mM NaCl. The N-terminal His-tag and DsbC fusion protein was removed by the cleavage with TEV protease (1:200 molar ratio) incubated at 4 °C for 24 h. Protein-containing fractions were pooled and concentrated on an Amicon 3 kDa MWCO spin concentrator (Merck Millipore). After another IMAC purification step using Ni-NTA resin, the flow-through was injected onto a reversed phase column (2 mL, Source 15RPC, Cytiva) with prior addition of acetonitrile and trifluoroacetic acid to achieve a concentration of 10% and 0.1%, respectively. Proteins were eluted with a linear gradient of 20% to 40% ACN containing 0.1% TFA. Pooled fractions were dried under vacuum in a centrifugal concentrator and aliquots stored at -80 °C. Reduced and non-reduced SDS–PAGE was used to analyse the purified proteins.

### Calcium imaging

Hypothalamic progenitors differentiated from the HUES9-POMC-GFP cell line were re-plated onto 35 mm x 10 mm polystyrene imaging dishes (Corning) coated with 1 µg/cm^2^ of iMatrix-511 (Takara Bio). Hypothalamic cultures were loaded with the Fluorescent Dye-Based Cal-590 following the manufacturers’ instructions (Stratech). Hanks’ Balanced Salt Solution Phenol Red free (HBSS; ThermoFisher scientific) with or without synaptic blockers were used as extracellular bath solutions. The dye loading, incubation and wash buffers were also prepared in HBSS. The synaptic blockers DL-AP5 100 µM, Picrotoxin 50 µM, CNQX 30 µM, and Strychnine 20 µM were added in all experiments where the responses to Semaglutide were measured. Experiments with ALKAL2 were performed in the absence of synaptic blocker. Stock solutions of 1 mM ALKAL2 (in PBS), 1 mM Semaglutide (in DMSO), and 1 mM ceritinib dihydrochloride (in DMSO) were stored at -20°C until the day of each experiment. Final concentrations of 300 nM ALKAL2 without synaptic blockers and 200 nM of Semaglutide with synaptic blockers were prepared in HBSS immediately before each experiment. For the Ceritinib pre-treatment experiment, 1 µM of Ceritinib dihydrochloride (MedChem Express) was added to hypothalamic cultures and incubated for 24 hours at 37°C in a humidified atmosphere of 5% CO2.

Imaging dishes were placed on an Olympus BX51WI Fixed Stage Upright Microscope (RRID: SCR_023069) and imaged using a 16-bit high-speed ORCA Flash4.0 LT plus digital sCMOS camera (RRID: SCR_021971). Neurons were identified by Cal-590 AM fluorescence using an excitation wavelength of 540 nm and an emission wavelength of 590 nm. Images were taken using a 20X objective and acquired at a frequency of 0.1 Hz (100 ms exposure/frame) using a CoolLED pE-300 white (RRID: SCR_021073) illumination system and HCImage (RRID: SCR_015041) software for acquisition. Neurons were perfused continuously at room temperature with HBSS solution (3 mL/min) using a gravity-driven perfusion system for the entire length of the experiment. Each experiment’s time course was as follows: After a variable washout period to ensure a stable baseline, cells were perfused with extracellular bath solution containing the tested drug (ALKAL2, 300 nM for 5 minutes; Semaglutide, 200 nM for 2 minutes). Following a second washout period (>10 minutes), neurons were stimulated with 50 mM KCl for two minutes. Fluorescence time course was recorded for the entire length of the experiment

Each fluorescence time course was exported to Microsoft Excel (RRID:SCR_016137), and the change in fluorescence intensity as a function of time was expressed as (F − F0)/F0) or ΔF/F0, where F is the total fluorescence and F0 is the baseline fluorescence. The extent of ALKAL2 or Semaglutide response was evaluated by comparing the change in the area under the curve (AUC) of the fluorescence time course before and after the perfusion with the agonist.

### Mouse purchase and housing conditions

All animal experiments were performed under the listed procedures detailed in UK Home Office-approved Project Licence PP7597478, and were approved by the University of Cambridge Animal Welfare and Ethics Review Board (AWERB) and adhered to 3Rs and ARRIVE guidelines. All mice used in this study were male C57BL/6J mice purchased from Charles Rivers Laboratories (Saffron Walden, UK) at an age of six to seven weeks. Upon arrival, mice were housed in groups of 4 unless specified otherwise in individually ventilated cages for a week to acclimatise them to the specific pathogen free facility and were given access to water and standard chow *ad libitum*. The animal facility was maintained on a standard 12-hour light/dark cycle (lights on at 0700h) and temperature and humidity was controlled at 22 ± 2°C and 55 ± 2% respectively.

### Drug effects on mouse body weight and body composition

For the acute injection experiment, standard laboratory chow was replaced with either a 60% high fat diet (HFD, D12492i, Research diets) or matched control (D12450Ji, Research diet). After 1 week on the modified diets, mice were given a single intraperitoneal injections of 10 mg/kg Ceritinib dihydrochloride (MedChem Express) or vehicle (dimethyl sulfoxide; Sigma Aldrich) that were dilution matched with an injection volume of 10 mL/kg between 1000h and 1100h. At 6- or 24-hours post-injection, body weights were recorded and mice were anesthetised with isoflurane. A terminal blood sample was collected from the heart in heparinised tubes (Greiner Bio-One) and centrifuged at 2000 x g for 10 min at 4°C. The resulting plasma was stored at -20°C for later biochemical analyses. The whole brain was rapidly removed and the hypothalamus was isolated, snap frozen on dry ice, and stored at -70°C for later qRT-PCR analysis.

For weight loss experiments over a two week period, standard laboratory chow was replaced with HFD for four weeks to generate a diet-induced obese (DIO) mice that were then maintained on HFD and injected intraperitoneally every 48 hours with candidate drugs or vehicle (DMSO; Sigma Aldrich) with an injection volume of 10 mL/kg. Drugs tested and concentrations given per injections were: 10 mg/kg Beta-lapachone (Bio-Techne), 10 mg/kg Ceritinib dihydrochloride, 40 μg/kg Semaglutide (MedChem Express), and 2 mg/kg Varenicline tartrate (Stratech). Body weight measurements and injections were performed between 10:00h and 11:00h to minimise potential circadian fluctuations. Across all treatment groups, no mice exceeded a limit of 20% reduction in body weight compared to vehicle controls. After two weeks of treatment (day 14), body composition analyses were performed using Echo MRI^TM^-100H Body Composition Analysers to estimate fat and lean mass. At day

15 of the experiment, one batch of mice treated with vehicle, Semaglutide, Ceritinib dihydrochloride, and Certinib + Semaglutide (n=4/group) were subjected to an intraperitoneal glucose tolerance test (ipGTT). At day 16 of the experiment, mice were then injected with one last dose of the allocated drug, and 2 hours later anesthetised with isoflurane. A terminal blood sample was collected in heparinised tubes (Greiner Bio-One) and centrifuged at 2000 x g for 10 min at 4°C. The resulting plasma was stored at -20°C for later biochemical analyses.

### ipGTT

Mice were fasted for 5 hours (maintaining free access to water) then subjected to the ipGTT with 2 g/kg body weight of glucose in normal saline. Tail blood was used to measure glucose concentration (mmol/L) using a standard glucometer (AlphaTRAK 2, Zoetis) at 0, 30, 60, 90, and 120 minutes.

### Drug effects on food intake and energy expenditure using metabolic cages

As described above, mice were group-housed on arrival and given standard laboratory chow during the acclimatisation period. They were then fed a 60% high fat diet (Research diets) and maintained on this diet for 3-6 weeks. Mice were then transferred and acclimatised to individual housing in metabolic cages (Promethion Metabolic Screening, Sable Systems) for five to seven days before any injections were made or measurements were taken. After the acclimatisation period, intraperitoneal injections (10 mL/kg injection volume) of vehicle (DMSO), 10 mg/kg Ceritinib dihydrochloride, 40 μg/kg Semaglutide, 20 mg/kg YNT-185 (Bio-Techne), or a combination of 10 mg/kg Ceritinib dihydrochloride and 40 μg/kg Semaglutide were performed every day at around 10:00h during which body weights were recorded. Each cage was equipped with water bottles and food hoppers connected to load cells (MM-1, Sable Systems International) for food and water intake monitoring. Mice had *ad libitum* access to food (60% high fat diet) and water throughout the study. In addition to food and water intake, energy expenditure was continuously recorded for 72 hours. On the day of tissue collection, mice were first injected with one last dose of the allocated drug and 2 hours later anaesthetised with isoflurane. The whole brain was rapidly removed and the hypothalamus was isolated, snapped frozen on dry ice, and immediately stored at -70°C for later bulk RNA sequencing and qRT-PCR analyses. Additionally, white epididymal adipose tissues were collected, snap frozen and stored at -70°C for later qRT-PCR analysis.

### Quantitative RT-PCR

hPSCs and hypothalamic progenitors were re-plated onto 12-well polystyrene plates (Corning) coated with Geltrex^TM^. Total RNA from cell pellets were extracted using the RNeasy Mini Kit (Qiagen) and treated with Deoxyribonuclease I (Qiagen) according to the manufacturer’s instructions. Extracted total RNA (0.25 - 0.75 µg) was reverse transcribed into cDNA using the iScript^TM^ reverse transcription supermix (Bio-Rad Laboratories). Target genes of interest were determined using the Taqman gene expression ‘assay-on-demand^TM^’ assays (Applied Biosystems, Foster City, CA) with FAM-labeled probes for target genes and beta actin (ACTB; Hs01060665_g1) and glyceraldehyde 3-phosphate dehydrogenase (GAPDH; Hs99999905_m1) as reference genes. The primer/probe targets used for assessing the differentiation of hypothalamic progenitors were DCX (Hs00167057_m1), ISL1 (Hs00158126_m1), NR5A1 (Hs00610436_m1), ASCL1 (Hs00269932_m1), NANOG (Hs02387400_g1), OTP (Hs01888165_s1), NES (Hs04187831_g1), and NKX2-1 (Hs00968940_m1). The primer/probe targets used for assessing the metabolic response to 24 hours of Ceritinib (1 µM) pre-treatment were GLP1R (Hs01379269_m1), LEPR (Hs00174497_m1), and INSR (Hs00961557_m1).

Total RNA from the mouse whole hypothalamus and epididymal white adipose tissue were extracted using the RNeasy Universal Plus Kit (Qiagen) according to the manufacturer’s instructions. Extracted total RNA (0.75 - 1 µg) was reverse transcribed into cDNA using the iScript^TM^ reverse transcription supermix. Target genes of interest in the hypothalamus were determined using the Taqman gene expression ‘assay-on-demand^TM^’ assays (Applied Biosystems, Foster City, CA) using FAM-labelled probes for target genes and *Actb* (Mm02619580_g1) and *Gapdh* (Hs99999905_m1) as reference genes. The primer/probe targets were *Agrp* (Mm00475829_g1), *Glp1r* (Mm00445292_m1), *Insr* (Mm01211875_m1), *Lepr* (Mm00440181_m1), *Mc4r* (Mm00457483_s1), and *Pomc* (Mm00435874_m1). The reactions were run on the QuantStudio™ 5 Real-Time PCR System in 384-well plates (Thermo Fisher Scientific). All expression assays were normalised to the geometric mean of the reference genes *GAPDH/Gapdh* and *ACTB/Actb*.

Targeted gene expression in the epididymal white adipose tissue was carried out using SYBR® Green PCR master mix (Thermo Fisher Scientific). cDNA was generated from 2 µg isolated RNA using M-MLV reverse transcriptase (Promega) and diluted 1:15 for use in 12 µL qPCR reactions run on the StepOnePlus™ system (Applied Biosystem). Expression values were normalised to the geometric mean of two reference genes, *36b4* and *B2m*. All SYBR forward and reverse primers can be found listed in the key resource table.

The fold change in expression of each target gene was calculated using the “ΔΔCt” method where Ct is the PCR threshold cycle, using the formula 2^-(ΔCt where ΔCt = (Ct (Target gene) – Ct (geometric mean of reference genes)).

### Biochemical measurements

All biochemical analyses were conducted at the Cambridge Core Biochemical Assay Laboratory (CBAL). Plasma concentrations of alanine transaminase (ALT), aspartate aminotransferase (AST), cholesterol, and C-reactive protein (CRP) were run automated on the Siemens Dimension EXL analyser. Plasma GDF-15 was performed as an in-house microtitre plate-based two-site electrochemiluminescence immunoassay using the MesoScale Discovery assay platform (MSD, Rockville, Maryland, USA). The GDF-15 antibodies and standards were purchased as a DuoSet (R&D Systems). A mouse leptin kit (MesoScale Discovery) was used to determine plasma leptin concentrations according to the manufacturer’s instructions.

### Bulk RNA sequencing

Total RNA from the whole hypothalamus of mice treated with vehicle or drugs was extracted using the RNeasy Universal Plus Kit (QIAGEN) according to the manufacturer’s instructions. RNA quality was assessed with the Agilent 2100 Bioanalyser system using the RNA 6000 Nano kit (Agilent Technologies) and the mean RIN number across all 16 samples was 8.4. The polyA stranded libraries were generated using the Novogene NGS Stranded RNA Library Prep Set (Novogene). In brief, messenger RNA was purified from total RNA using poly-T oligo-attached magnetic beads. After fragmentation, the first strand cDNA was synthesised using random hexamer primers. Then the second strand cDNA was synthesised using dUTP instead of dTTP followed by end repair, A-tailing, adapter ligation, size selection, USER enzyme digestion, amplification, and purification. The resulting cDNA library was analysed by Qubit and real-time PCR for quantification and bioanalyzer for size distribution detection. Quantified libraries were pooled and sequenced on an Illumina platform to generate 150bp paired-end reads.

### Bulk RNA sequencing analysis

Raw bulk RNA sequencing libraries were pre-processed with TrimGalore (v0.6.7) (Krueger et al. 2021) for adapter removal and filtering of low-quality reads. Reads were aligned using STAR (v2.7.9a) (Dobin et al. 2013) to either human (GRCh38, Ensembl release 98) or mouse (GRCm38, Ensembl release 98) reference genome, and then quantified using featureCounts (v2.0.0) (Liao, Smyth, and Shi 2014) against the human (GRCh38, gencode v32) or mouse annotation (GRCm38, gencode vM23), respectively. Qualimap (v2.2.2d) (Okonechnikov, Conesa, and García-Alcalde 2015) was used to generate QC metrics of aligned reads for evaluating sample library quality, together with QC metrics produced by STAR and featureCounts. Filtering of lowly expressed genes and size factor for normalisation of the bulk gene counts was performed using the edgeR package, as described previously. Principal component analysis and expression correlation analysis were then performed, followed by estimation of both negative binomial dispersions and quasi-likelihood dispersion using the edgeR package. Differential expression testing was performed using a quasi-likelihood F-test to identify genes which are differentially expressed between different treatment groups, with significance threshold set to absolute log2(fold-change) > 0 and FDR < 0.1. Finally, gene set enrichment analysis of the differentially expressed genes were performed using the gost function from the g:Profiler R package (Raudvere et al. 2019) with the expressed gene list set as the custom background and analytical-adjusted p-value threshold set to the default 0.05. Gene sets were then filtered for sets containing 5-500 genes to avoid higher-level gene sets.

### Peptidomics

Whole hypothalamus collected from a cohort of mice treated with 5 days of vehicle or drugs were homogenised in 6M guanidine hydrochloride, and extracted using acetonitrile precipitation and solid phase extraction as previously described (Kirwan et al. 2018). The extracted peptides were reduced and alkylated and then analysed by nanoflow LC-MS/MA using a ThermoScientific Ultimate 3000 nano LC system coupled to a Q-Exactive Plus Orbitrap mass spectrometer (ThermoScientific). Relative peptide quantification was performed using Qualbrowser (ThermoScientific) for Agrp and desacetyl-α-MSH.

### Statistical analyses

Graphing and statistical analysis were performed using GraphPad Prism (version 7.02; RRID: SCR_002798). For all one-way analyses of variance (ANOVA), data were first analysed for normality using the Brown–Forsythe test. Non-parametric Kruskal-Wallis ANOVA with Dunn’s multiple comparisons test was used for data sets with significantly different standard deviations. One-way ANOVA with Dunnett’s multiple comparison test was used to compare the quantified percentage of MAP2^+^ cells expressing POMC across different culture conditions. One-way ANOVA with Tukey’s post hoc test was used to compare the intrinsic membrane properties of neurons cultured in SJ versus NB media. A non-parametric paired t-test was used to determine whether the responses of ALKAL2 or Semaglutide significantly differed from the baselines. Two-way ANOVA with Bonferroni’s multiple comparisons test was used to compare the effects of candidate drugs on weight changes over time. One-way ANOVA with Dunnett’s multiple comparison test was used to compare changes in EchoMRI and biochemical parameters between different drug treatment groups. The effects of drugs on energy expenditure and food intake was analysed by both one-way ANOVA with Dunnett’s multiple comparison test and one-way analysis of covariance (ANCOVA). One-way ANOVA with Dunnett’s multiple comparison test was used to compare Sem, Cer, and Com to Veh treatment groups from quantitative RT-PCR and peptidomics analyses. Unpaired t test was used to compare between Veh and Cer in the acute injection (24 hour treatment) and *in vitro* Ceritinib stimulation experiment. A non-parametric unpaired t-test was used to determine whether the responses to Semaglutide significantly differed between neurons pre-treated for 24 hours with vehicle or 1 µM Ceritinib. Results were presented as mean with 95% CI for all calcium imaging experiments and mean ± standard error of the mean (SEM) for all other experiments. After adjusting for multiple comparisons, P values less than 0.05 were considered as statistically significant.

### Data, code, and materials availability

Upon publication in a peer-reviewed journal, sequencing data and processed data of scRNAseq and bulk RNAseq data will be made available on ENA, and code used for analysis and generating figures will be made available in GitHub. Reagents will be made available upon request from the lead author (F.T.M.).

## Author contributions

**H.-J.C.C:** Conceptualisation, Methodology, Investigation, Formal analysis, Visualisation, Writing – Original Draft, Writing – Review & Editing. **A.Y:** Conceptualisation, Methodology, Software, Investigation, Formal analysis, Visualisation, Writing – Original Draft, Writing – Review & Editing. **S.M:** Methodology, Investigation, Formal analysis, Visualisation, Writing – Review & Editing. **I.M:** Methodology, Investigation, Writing – Review & Editing. **O.C:** Investigation. **K.K**: Methodology, Formal analysis, Writing – Review & Editing. **C.R:** Investigation. **N.S:** Investigation. **J.W.E.S:** Project administration. **V.P:** Investigation. **P.K:** Investigation. **S.A:** Investigation. **T.T:** Resources, Investigation. **M.H.C.F:** Resources, Investigation. **C.A.C:** Resources. **L.Q:** Investigation, Writing – Review & Editing. **M.H:** Resources, Writing – Review & Editing. **M.R.L:** Investigation, Writing – Review & Editing. **J.R.B.P:** Methodology, Investigation, Writing – Review & Editing. **J.C.M:** Conceptualisation, Software, Investigation, Resources, Supervision, Writing – Review & Editing. **F.T.M:** Conceptualisation, Investigation, Resources, Supervision, Writing – Original Draft, Writing – Review & Editing, Project administration, Funding acquisition. All authors have reviewed the manuscript and approve of its content. Co-first authors (**H.-J.C.C** and **A.Y**) contributed equally to this manuscript and all authors agree that they can indicate their equal contribution and re-order the list of co-first authors in their own publication records.

## Acknowledgements

We thank Davide Chiarugi and Katherine Lawler for their discussions and advice that helped shape this work. A.Y. was supported by an EMBL-EBI/Cambridge Computational Biomedical Postdoctoral Fellowship (EBPOD). V.P. is supported by a UK Regenerative Medicine Platform grant from the Medical Research Council (MR/R015724/1). L.Q. is grateful for a Walter-Benjamin Fellowship by the German Research Council. F.T.M. is a New York Stem Cell Foundation - Robertson Investigator (NYSCF-R-156) and is supported by the Wellcome Trust and Royal Society (211221/Z/18/Z) and a Ben Barres Early Career Acceleration Award from the Chan Zuckerberg Initiative’s Neurodegeneration Challenge Network (CZI NDCN 191942). J.C.M. acknowledges core funding from the European Molecular Biology Laboratory and from Cancer Research UK (C9545/A29580). We thank Professor Anne White for kindly providing the A1H5 antibody, and thank Amit Chouhan and Ann Cloos for their assistance with tissue culture. We appreciate the assistance from Greg Strachnan and the MRC Metabolic Diseases Unit Imaging Core Facility with imaging and analysis on the Operphenix. We would like to thank Katarzyna Kania and the staff at the Genomics Core Facility of the Cancer Research UK Cambridge Institute for assistance with single-cell library preparation and sequencing in addition to Davide Chiarugi and other members of the Bioinformatics and Biostatistics (Bio2) core facility for their help on data management. We would like to thank Peter Barker, Keith Burling and other members of the Cambridge Biochemical Assay Laboratory (CBAL) for their assistance running mouse plasma samples. We sincerely thank Richard Kay and the IMS Peptidomics and Proteomic Core Facility for running the tissue peptide analysis. We are extremely grateful to the assistance of Marcella Ma and the IMS Genomics and Transcriptomics Core Facility, as well as Alix Schwiening and the Disease Model Core (DMC), part of the MRC Metabolic Diseases Unit (MC_UU_00014/5) both of which are funded by the UK Medical Research Council (MRC) Metabolic Disease Unit (MRC_MC_UU_00014/5), Wellcome Trust Major Award (208363/Z/17/Z), and a Wellcome DRP. For the purpose of open access, the authors have applied a CC-BY public copyright licence to any Author Accepted Manuscript version arising from this submission.

---

**Figure S1.**
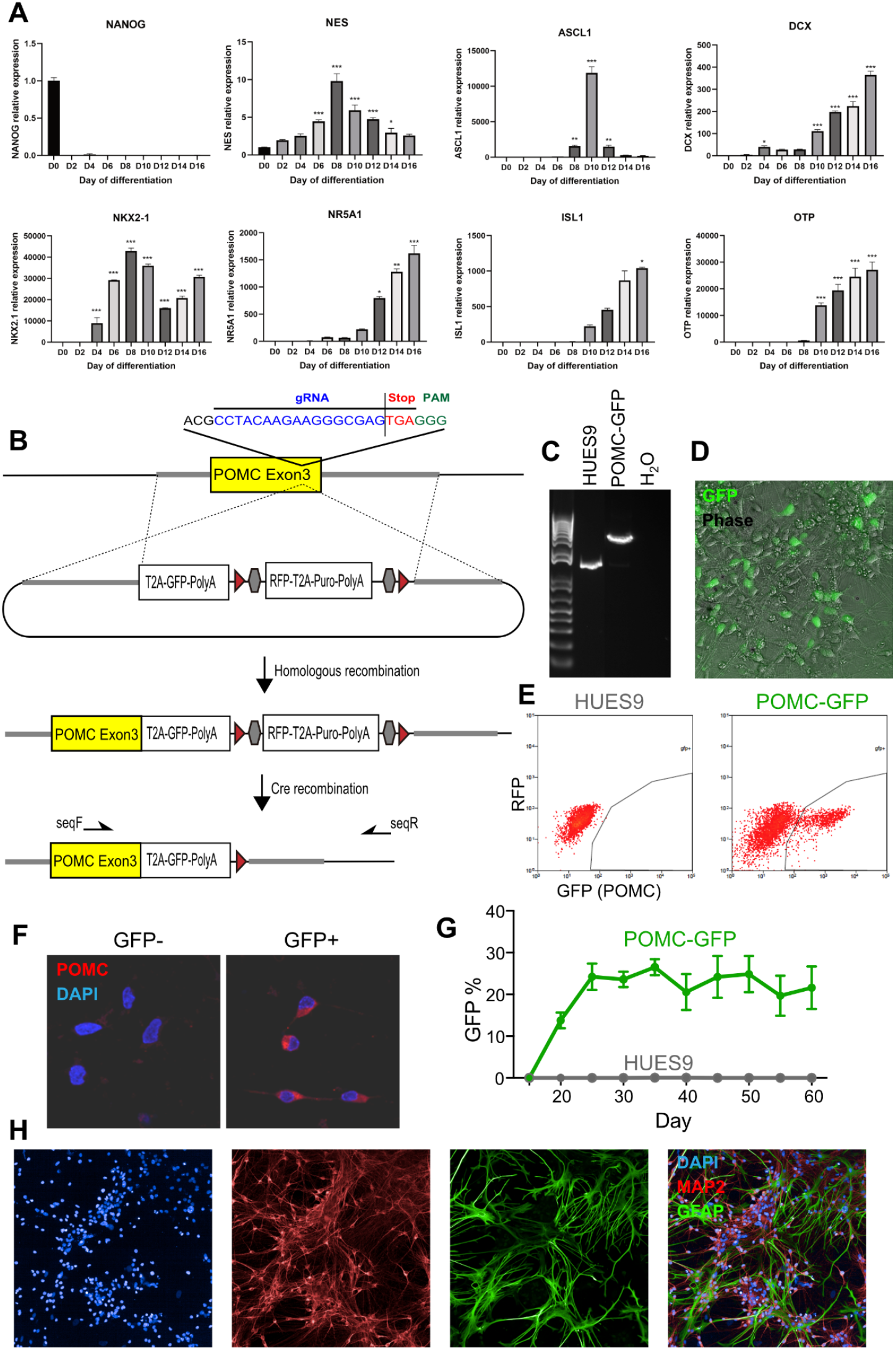
Generation of a POMC-GFP cell line, and astrocyte co-culture. **A)** Gene expression time course during hypothalamic differentiation in the KOLF2.1J cell line. **B)** Schematic used for CRISPR/Cas9-based knock-in of a T2A-GFP-polyA cassette to replace the TGA Stop codon of the endogenous human *POMC* gene in the HUES9 human embryonic stem cell line. **C)** PCR confirming the correct 5’ targeting of the *POMC* gene using primers indicated in (B). **D)** Representative photomicrograph of a differentiated culture of the POMC-GFP reporter gene, with GFP channel merged with phase contrast channel. **E)** FACS plots from the parental and gene edited cell line after hypothalamic differentiation. **F)** FAC sorting of GFP- and GFP+ cell fractions followed by culture and immunostaining confirms that 94±3% of GFP+ cells were immunoreactive for POMC whereas 1±1% of GFP-cells were (n=3 differentiations). **G)** Time course of GFP+ cell generation from HUES9 or POMC-GFP cell lines differentiated to hypothalamic neurons, as quantified by FACS (n=3 replicates). **H)** Representative photomicrographs of human hypothalamic neurons (MAP2, red) co-cultured with mouse primary astrocytes (GFAP, green).

**Figure S2.**
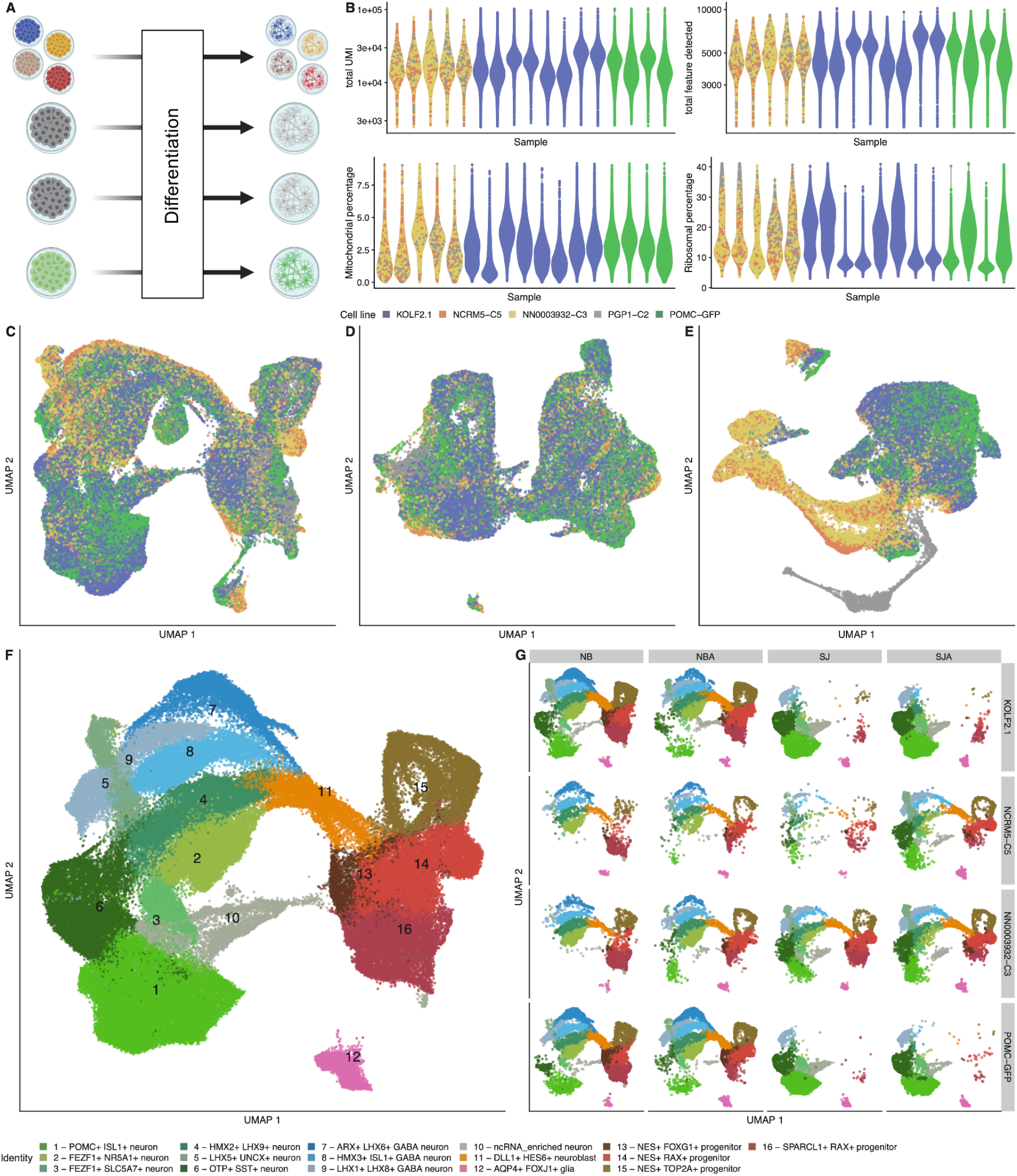
Hypothalamic differentiation is efficient regardless of genetic background or differentiation batch. **A)** Experimental schematic showing pooled or individual cell differentiation in different batches. **B)** Quality control of sequencing data shows abundant UMIs and features per cell across samples and cell lines, with consistently low mitochondrial and modest ribosomal gene expression. **C-E)** UMAP representation of single cell data from all cell lines and culture conditions (C), reveals a relatively even contribution of cell lines to each cell cluster in N2B27-matured (D) and SJ-matured (E) cultures, with the exception of of cell line PGP1. **F)** After removing line PGP1 and re-clustering data, neuronal cell sub-populations (shades of green and blue) are clearly distinguishable from neuroblasts (orange), and progenitors and glia (shades of red and green), whose gene expression patterns identify them as predominantly resembling the anterior ventral hypothalamus including the arcuate nucleus and ventromedial hypothalamus. **G)** All four genetically distinct cell lines (rows) contributed relatively evenly to all cell clusters, whereas cells matured in N2B27-based maturation medium (columns NB and NBA) contributed to more progenitors and fewer neurons than those matured in Synaptojuice-based medium (columns SJ and SJA).

**Figure S3.**
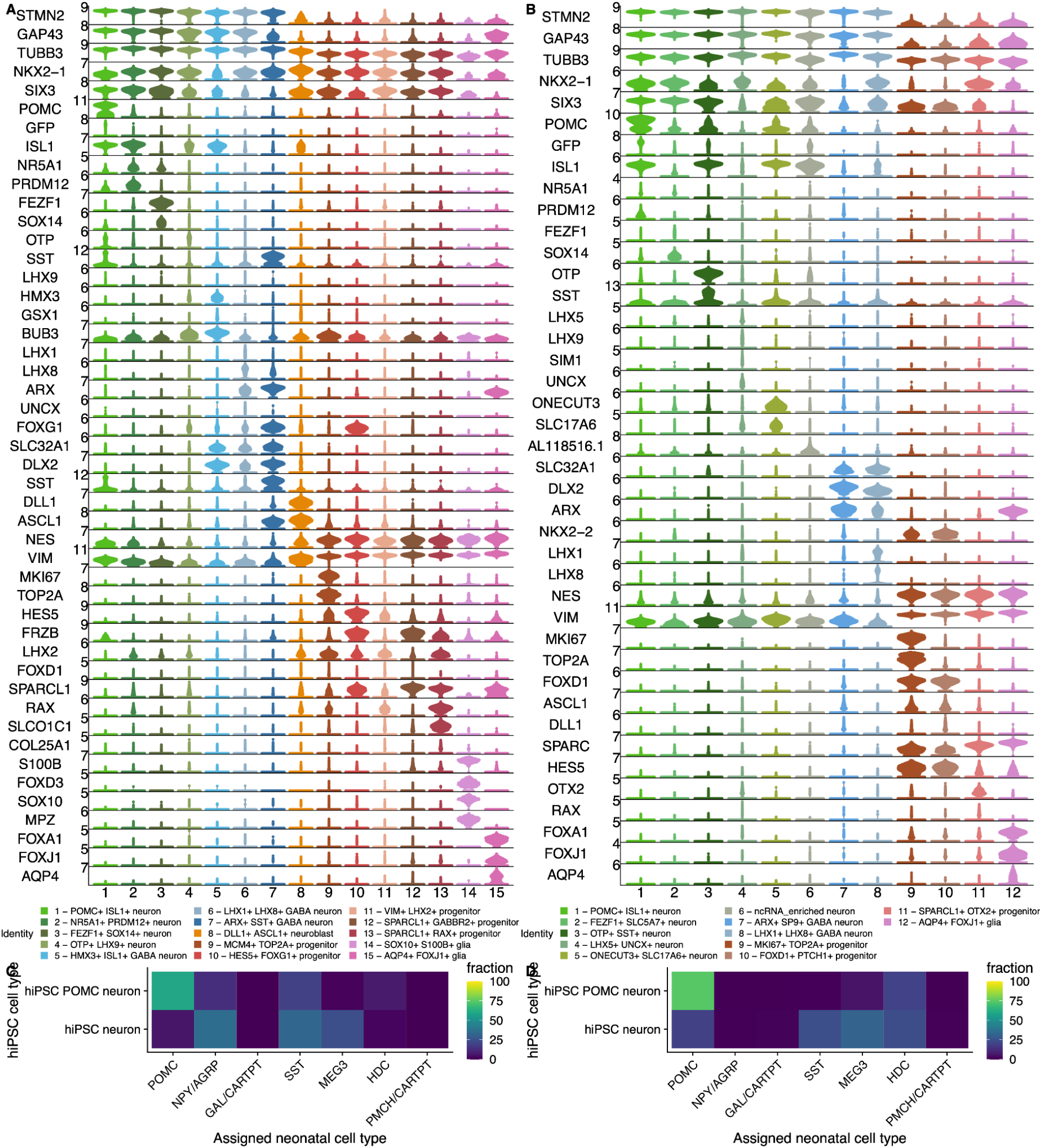
hPSC-derived hypothalamic cell types resemble their counterparts *in vivo*. **A, B)** Genes enriched in clusters of cells matured in N2B27-based medium (A) or Synaptojuice-based medium indicate that hPSC-derived cells have spatial identities indicative of predominantly hypothalamic identity as described in Results. **C,D)** Unbiased label transfer analysis indicates that the vast majority of hPSC-derived cells annotated as POMC neurons are also annotated as POMC neurons in from the neonatal human arcuate nucleus.

**Figure S4.**
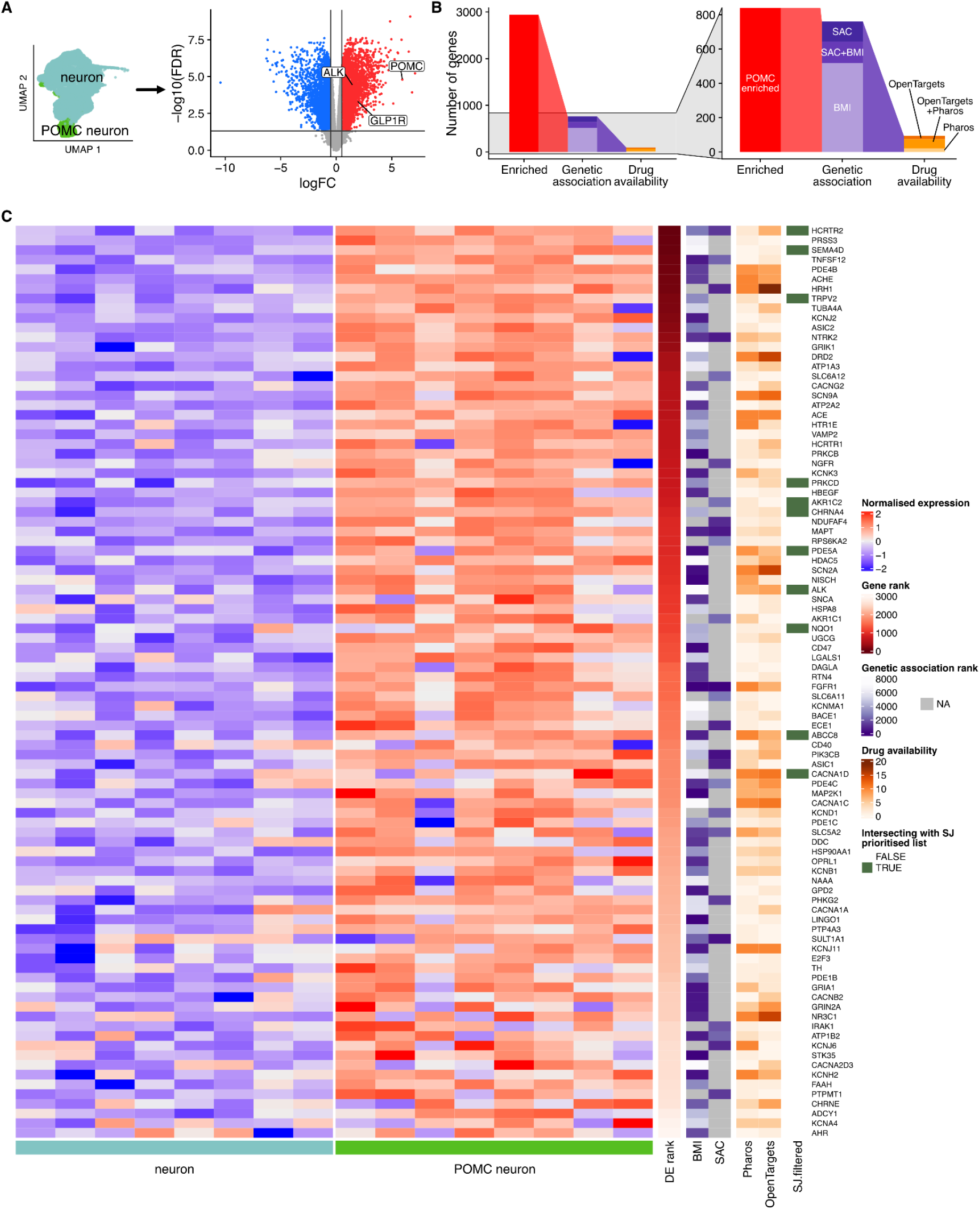
Identification of candidate drugs in POMC neurons matured in N2B27 media. **A)** Schematic of POMC neuron and non-POMC neuron classification and volcano plot resulting from differential expression analysis. **B)** Waterfall plot of scheme used to identify candidate genes that were enriched in NB-matured POMC neurons, genetically associated with obesity, and targeted by drugs approved for human use. **C)** Summary of candidate genes following the scheme used in Figure 3, and with genes identified in hPOMC neurons grown in both media types (including ALK) indicated in green

**Figure S5.**
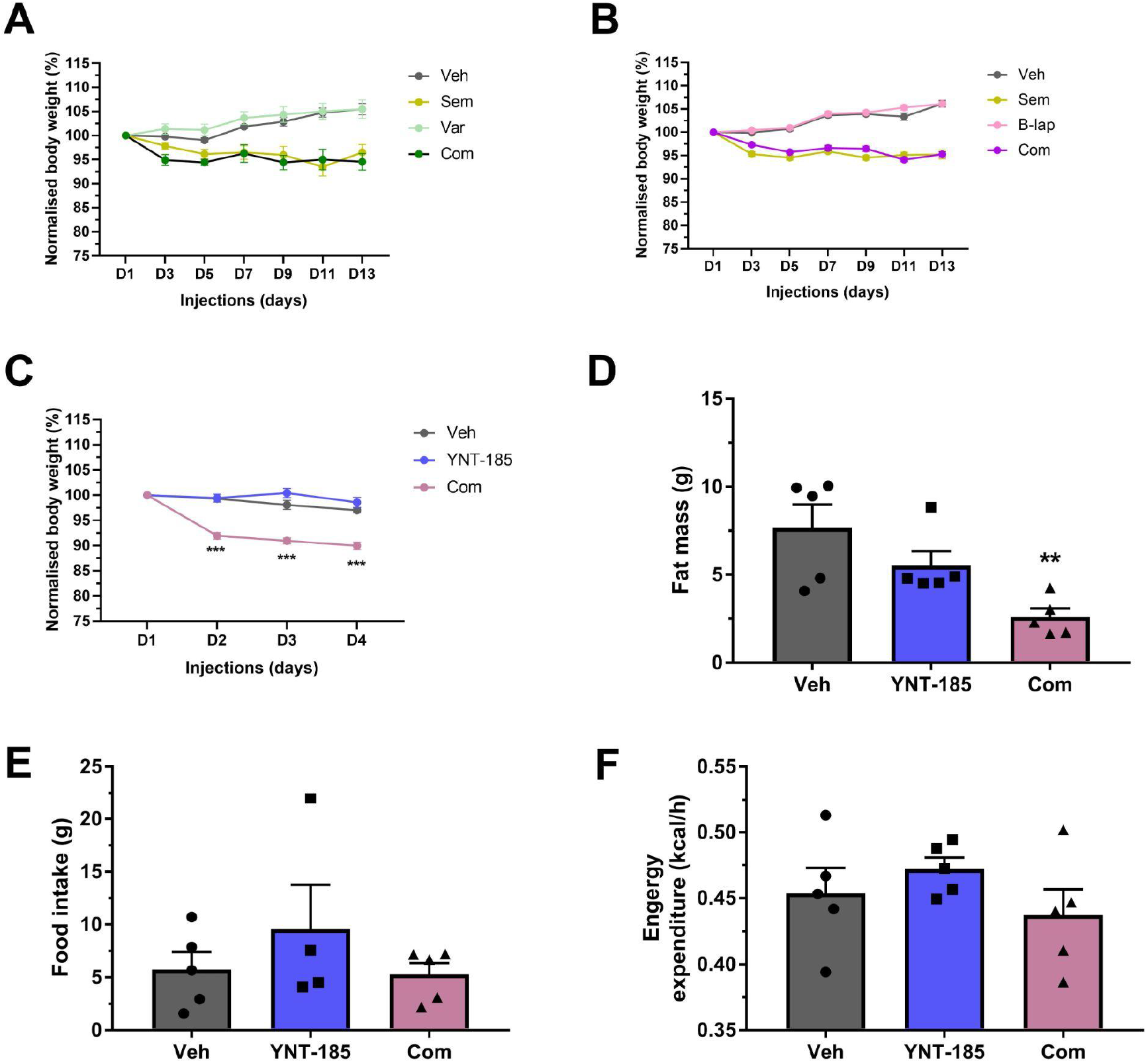
Summary of results from other drugs tested for weight loss effects in DIO mouse models. While Semaglutide (Sem) was effective, no significant weight reducing effects were observed with candidate drugs alone, nor were additive effects seen in combinatorial (Com) treatment. Drugs used included: **(A)** the cholinergic receptor nicotinic alpha 4 subunit (*CHRNA4*) agonist varenicline (Var; n=4/group), **(B)** the NAD(P)H quinone dehydrogenase 1 (*NQO1*) substrate beta-lapachone (B-lap; n=8/group), or **(C)** the hypocretin receptor type 2 (*HCRTR2*) agonist YNT-185 (n=5/group), which also did not affect fat mass **(D)**, food intake **(E)**, or energy expenditure **(F)**. Data represent mean ± SEM; **p<0.01 and ***p<0.001.

## References

Ahmed, Mansoor, Navjot Kaur, Qianni Cheng, Marya Shanabrough, Evgenii O. Tretiakov, Tibor Harkany, Tamas L. Horvath, and Joseph Schlessinger. 2022. “A Hypothalamic Pathway for Augmentor α-Controlled Body Weight Regulation.” Proceedings of the National Academy of Sciences of the United States of America 119 (16): e2200476119.

Alvarez-Bolado, Gonzalo, Valery Grinevich, and Luis Puelles. 2015. Development of the Hypothalamus. Frontiers Media SA.

Aponte, Yexica, Deniz Atasoy, and Scott M. Sternson. 2011. “AGRP Neurons Are Sufficient to Orchestrate Feeding Behavior Rapidly and without Training.” Nature Neuroscience 14 (3): 351–55.

Bais, Abha S., and Dennis Kostka. 2019. “Scds: Computational Annotation of Doublets in Single-Cell RNA Sequencing Data.” Bioinformatics 36 (4): 1150–58.

Balland, Eglantine, Julie Dam, Fanny Langlet, Emilie Caron, Sophie Steculorum, Andrea Messina, S. Rasika, et al. 2014. “Hypothalamic Tanycytes Are an ERK-Gated Conduit for Leptin into the Brain.” Cell Metabolism 19 (2): 293–301.

Bedont, Joseph L., Tara A. LeGates, Emily A. Slat, Mardi S. Byerly, Hong Wang, Jianfei Hu, Alan C. Rupp, et al. 2014. “Lhx1 Controls Terminal Differentiation and Circadian Function of the Suprachiasmatic Nucleus.” Cell Reports 7 (3): 609–22.

Bedont, Joseph L., Elizabeth A. Newman, and Seth Blackshaw. 2015. “Patterning, Specification, and Differentiation in the Developing Hypothalamus.” Wiley Interdisciplinary Reviews. Developmental Biology 4 (5): 445–68.

Bilican, Bilada, Matthew R. Livesey, Ghazal Haghi, Jing Qiu, Karen Burr, Rick Siller, Giles E. Hardingham, David J. A. Wyllie, and Siddharthan Chandran. 2014. “Physiological Normoxia and Absence of EGF Is Required for the Long-Term Propagation of Anterior Neural Precursors from Human Pluripotent Cells.” PloS One 9 (1): e85932.

Bilsland, James G., Alan Wheeldon, Andrew Mead, Petr Znamenskiy, Sarah Almond, Kerry A. Waters, Matthew Thakur, et al. 2008. “Behavioral and Neurochemical Alterations in Mice Deficient in Anaplastic Lymphoma Kinase Suggest Therapeutic Potential for Psychiatric Indications.” Neuropsychopharmacology: Official Publication of the American College of Neuropsychopharmacology 33 (3): 685–700.

Burmeister, Melissa A., Jennifer E. Ayala, Hannah Smouse, Adriana Landivar-Rocha, Jacob D. Brown, Daniel J. Drucker, Doris A. Stoffers, Darleen A. Sandoval, Randy J. Seeley, and Julio E. Ayala. 2017. “The Hypothalamic Glucagon-Like Peptide 1 Receptor Is Sufficient but Not Necessary for the Regulation of Energy Balance and Glucose Homeostasis in Mice.” Diabetes 66 (2): 372–84.

Bycroft, Clare, Colin Freeman, Desislava Petkova, Gavin Band, Lloyd T. Elliott, Kevin Sharp, Allan Motyer, et al. 2018. “The UK Biobank Resource with Deep Phenotyping and Genomic Data.” Nature 562 (7726): 203–9.

Challis, B. G., A. P. Coll, G. S. H. Yeo, S. B. Pinnock, S. L. Dickson, R. R. Thresher, J. Dixon, et al. 2004. “Mice Lacking pro-Opiomelanocortin Are Sensitive to High-Fat Feeding but Respond Normally to the Acute Anorectic Effects of Peptide-YY(3-36).” Proceedings of the National Academy of Sciences of the United States of America 101 (13): 4695–4700.

Chen, Hsiao-Jou Cortina, Simone Mazzaferro, Tian Tian, Iman Mali, and Florian T. Merkle. 2023. “Differentiation, Transcriptomic Profiling, and Calcium Imaging of Human Hypothalamic Neurons.” Current Protocols 3 (6): e786.

Chen, Renchao, Xiaoji Wu, Lan Jiang, and Yi Zhang. 2017. “Single-Cell RNA-Seq Reveals Hypothalamic Cell Diversity.” Cell Reports 18 (13): 3227–41.

Chen, Yunshun, Aaron T. L. Lun, and Gordon K. Smyth. 2016. “From Reads to Genes to Pathways: Differential Expression Analysis of RNA-Seq Experiments Using Rsubread and the edgeR Quasi-Likelihood Pipeline.” F1000Research 5 (1438): 1438.

Chow, Laura Q. M., Fabrice Barlesi, Erin M. Bertino, Martin J. van den Bent, Heather A. Wakelee, Patrick Y. Wen, Chao-Hua Chiu, et al. 2022. “ASCEND-7: Efficacy and Safety of Ceritinib Treatment in Patients with ALK-Positive Non-Small Cell Lung Cancer Metastatic to the Brain And/or Leptomeninges.” Clinical Cancer Research: An Official Journal of the American Association for Cancer Research 28 (12): 2506–16.

Csárdi, Gábor, Tamás Nepusz, Kirill Müller, Szabolcs Horvát, Vincent Traag, Fabio Zanini, and Daniel Noom. 2023. “Igraph for R: R Interface of the Igraph Library for Graph Theory and Network Analysis,” June. https://doi.org/10.5281/zenodo.8046777.

Czerny, Christian Carl, Anett Borschel, Mingfang Cai, Madeline Otto, and Sigrid Hoyer-Fender. 2022. “FOXA1 Is a Transcriptional Activator of Odf2/Cenexin and Regulates Primary Ciliation.” Scientific Reports 12 (1): 21468.

Dhillon, Sohita, and Madeleine Clark. 2014. “Ceritinib: First Global Approval.” Drugs 74 (11): 1285–91.

Dobin, Alexander, Carrie A. Davis, Felix Schlesinger, Jorg Drenkow, Chris Zaleski, Sonali Jha, Philippe Batut, Mark Chaisson, and Thomas R. Gingeras. 2013. “STAR: Ultrafast Universal RNA-Seq Aligner.” Bioinformatics 29 (1): 15–21.

Drucker, Daniel J. 2018. “Mechanisms of Action and Therapeutic Application of Glucagon-like Peptide-1.” Cell Metabolism 27 (4): 740–56.

Elmentaite, Rasa, Natsuhiko Kumasaka, Kenny Roberts, Aaron Fleming, Emma Dann, Hamish W. King, Vitalii Kleshchevnikov, et al. 2021. “Cells of the Human Intestinal Tract Mapped across Space and Time.” Nature 597 (7875): 250–55.

Felix, Janine F., Jonathan P. Bradfield, Claire Monnereau, Ralf J. P. van der Valk, Evie Stergiakouli, Alessandra Chesi, Romy Gaillard, et al. 2016. “Genome-Wide Association Analysis Identifies Three New Susceptibility Loci for Childhood Body Mass Index.” Human Molecular Genetics 25 (2): 389–403.

Gabery, Sanaz, Casper G. Salinas, Sarah J. Paulsen, Jonas Ahnfelt-Rønne, Tomas Alanentalo, Arian F. Baquero, Stephen T. Buckley, et al. 2020. “Semaglutide Lowers Body Weight in Rodents via Distributed Neural Pathways.” JCI Insight 5 (6). https://doi.org/10.1172/jci.insight.133429.

Gelman, Diego M., Francisco J. Martini, Sandrina Nóbrega-Pereira, Alessandra Pierani, Nicoletta Kessaris, and Oscar Marín. 2009. “The Embryonic Preoptic Area Is a Novel Source of Cortical GABAergic Interneurons.” The Journal of Neuroscience: The Official Journal of the Society for Neuroscience 29 (29): 9380–89.

Giambartolomei, Claudia, Damjan Vukcevic, Eric E. Schadt, Lude Franke, Aroon D. Hingorani, Chris Wallace, and Vincent Plagnol. 2014. “Bayesian Test for Colocalisation between Pairs of Genetic Association Studies Using Summary Statistics.” PLoS Genetics 10 (5): e1004383.

Gribble, Fiona M., and Frank Reimann. 2019. “Function and Mechanisms of Enteroendocrine Cells and Gut Hormones in Metabolism.” Nature Reviews. Endocrinology 15 (4): 226–37.

GTEx Consortium. 2015. “Human Genomics. The Genotype-Tissue Expression (GTEx) Pilot Analysis: Multitissue Gene Regulation in Humans.” Science 348 (6235): 648–60.

Guan, Jikui, Ganesh Umapathy, Yasuo Yamazaki, Georg Wolfstetter, Patricia Mendoza, Kathrin Pfeifer, Ateequrrahman Mohammed, et al. 2015. “FAM150A and FAM150B Are Activating Ligands for Anaplastic Lymphoma Kinase.” eLife 4 (September): e09811.

Guillemot, F., L. C. Lo, J. E. Johnson, A. Auerbach, D. J. Anderson, and A. L. Joyner. 1993. “Mammalian Achaete-Scute Homolog 1 Is Required for the Early Development of Olfactory and Autonomic Neurons.” Cell 75 (3): 463–76.

Haghverdi, Laleh, Aaron T. L. Lun, Michael D. Morgan, and John C. Marioni. 2018. “Batch Effects in Single-Cell RNA-Sequencing Data Are Corrected by Matching Mutual Nearest Neighbors.” Nature Biotechnology 36 (5): 421–27.

Hallberg, B., and R. H. Palmer. 2016. “The Role of the ALK Receptor in Cancer Biology.” Annals of Oncology: Official Journal of the European Society for Medical Oncology / ESMO 27 Suppl 3 (September): iii4–15.

Helgeland, Øyvind, Marc Vaudel, Pol Sole-Navais, Christopher Flatley, Julius Juodakis, Jonas Bacelis, Ingvild L. Koløen, et al. 2022. “Characterization of the Genetic Architecture of Infant and Early Childhood Body Mass Index.” Nature Metabolism 4 (3): 344–58.

Herb, Brian R., Hannah J. Glover, Aparna Bhaduri, Carlo Colantuoni, Tracy L. Bale, Kimberly Siletti, Sten Linnarsson, et al. 2022. “Single-Cell Genomics Reveals Region-Specific Developmental Trajectories Underlying Neuronal Diversity in the Human Hypothalamus.” bioRxiv. https://doi.org/10.1101/2021.07.20.453090.

Hodge, Rebecca D., Trygve E. Bakken, Jeremy A. Miller, Kimberly A. Smith, Eliza R. Barkan, Lucas T. Graybuck, Jennie L. Close, et al. 2019. “Conserved Cell Types with Divergent Features in Human versus Mouse Cortex.” Nature 573 (7772): 61–68.

Huang, Wei-Kai, Samuel Zheng Hao Wong, Sarshan R. Pather, Phuong T. T. Nguyen, Feng Zhang, Daniel Y. Zhang, Zhijian Zhang, et al. 2021. “Generation of Hypothalamic Arcuate Organoids from Human Induced Pluripotent Stem Cells.” Cell Stem Cell 28 (9): 1657–70.e10.

Huang, Yuanhua, Davis J. McCarthy, and Oliver Stegle. 2019. “Vireo: Bayesian Demultiplexing of Pooled Single-Cell RNA-Seq Data without Genotype Reference.” Genome Biology 20 (1): 1–12.

Hu, Jia Sheng, Daniel Vogt, Magnus Sandberg, and John L. Rubenstein. 2017. “Cortical Interneuron Development: A Tale of Time and Space.” Development 144 (21): 3867–78.

Iwahara, T., J. Fujimoto, D. Wen, R. Cupples, N. Bucay, T. Arakawa, S. Mori, B. Ratzkin, and T. Yamamoto. 1997. “Molecular Characterization of ALK, a Receptor Tyrosine Kinase Expressed Specifically in the Nervous System.” Oncogene 14 (4): 439–49.

Jais, Alexander, and Jens C. Brüning. 2022. “Arcuate Nucleus-Dependent Regulation of Metabolism-Pathways to Obesity and Diabetes Mellitus.” Endocrine Reviews 43 (2): 314–28.

Kaminow, Benjamin, Dinar Yunusov, and Alexander Dobin. 2021. “STARsolo: Accurate, Fast and Versatile Mapping/quantification of Single-Cell and Single-Nucleus RNA-Seq Data.” bioRxiv. https://doi.org/10.1101/2021.05.05.442755.

Kemp, Paul J., David J. Rushton, Polina L. Yarova, Christian Schnell, Charlene Geater, Jane M. Hancock, Annalena Wieland, et al. 2016. “Improving and Accelerating the Differentiation and Functional Maturation of Human Stem Cell-Derived Neurons: Role of Extracellular Calcium and GABA.” The Journal of Physiology 594 (22): 6583–94.

Kentistou, Katherine A., Lena R. Kaisinger, Stasa Stankovic, Marc Vaudel, Edson M. de Oliveira, Andrea Messina, Robin G. Walters, et al. 2023. “Understanding the Genetic Complexity of Puberty Timing across the Allele Frequency Spectrum.” medRxiv. https://doi.org/10.1101/2023.06.14.23291322.

Kim, Dong Won, Parris Whitney Washington, Zoe Qianyi Wang, Sonia Hao Lin, Changyu Sun, Basma Taleb Ismail, Hong Wang, Lizhi Jiang, and Seth Blackshaw. 2020. “The Cellular and Molecular Landscape of Hypothalamic Patterning and Differentiation from Embryonic to Late Postnatal Development.” Nature Communications 11 (1): 4360.

Kirwan, Peter, Magdalena Jura, and Florian T. Merkle. 2017. “Generation and Characterization of Functional Human Hypothalamic Neurons.” Current Protocols in Neuroscience */ Editorial Board,* Jacqueline N. Crawley… [et Al.] 81 (October): 3.33.1–3.33.24.

Kirwan, Peter, Richard G. Kay, Bas Brouwers, Vicente Herranz-Pérez, Magdalena Jura, Pierre Larraufie, Julie Jerber, et al. 2018. “Quantitative Mass Spectrometry for Human Melanocortin Peptides in Vitro and in Vivo Suggests Prominent Roles for β-MSH and Desacetyl α-MSH in Energy Homeostasis.” Molecular Metabolism 17 (November): 82–97.

Kort, Anita, Rolf W. Sparidans, Els Wagenaar, Jos H. Beijnen, and Alfred H. Schinkel. 2015. “Brain Accumulation of the EML4-ALK Inhibitor Ceritinib Is Restricted by P-Glycoprotein (P-GP/ABCB1) and Breast Cancer Resistance Protein (BCRP/ABCG2).” Pharmacological Research: The Official Journal of the Italian Pharmacological Society 102 (December): 200–207.

Krueger, Felix, Frankie James, Phil Ewels, Ebrahim Afyounian, and Benjamin Schuster-Boeckler. 2021. “FelixKrueger/TrimGalore: v0.6.7 - DOI via Zenodo,” July. https://doi.org/10.5281/zenodo.5127899.

Kurrasch, Deborah M., Clement C. Cheung, Florence Y. Lee, Phu V. Tran, Kenji Hata, and Holly A. Ingraham. 2007. “The Neonatal Ventromedial Hypothalamus Transcriptome Reveals Novel Markers with Spatially Distinct Patterning.” The Journal of Neuroscience: The Official Journal of the Society for Neuroscience 27 (50): 13624–34.

Lee, Bora, Seunghee Lee, Soo-Kyung Lee, and Jae W. Lee. 2016. “The LIM-Homeobox Transcription Factor Isl1 Plays Crucial Roles in the Development of Multiple Arcuate Nucleus Neurons.” Development 143 (20): 3763–73.

Leeuw, Christiaan A. de, Joris M. Mooij, Tom Heskes, and Danielle Posthuma. 2015. “MAGMA: Generalized Gene-Set Analysis of GWAS Data.” PLoS Computational Biology 11 (4): e1004219.

Lee, Yung Seng, Ben G. Challis, Darren A. Thompson, Giles S. H. Yeo, Julia M. Keogh, Michael E. Madonna, Vicki Wraight, et al. 2006. “A POMC Variant Implicates Beta-Melanocyte-Stimulating Hormone in the Control of Human Energy Balance.” Cell Metabolism 3 (2): 135–40.

Lendahl, U., L. B. Zimmerman, and R. D. McKay. 1990. “CNS Stem Cells Express a New Class of Intermediate Filament Protein.” Cell 60 (4): 585–95.

Liao, Yang, Gordon K. Smyth, and Wei Shi. 2014. “featureCounts: An Efficient General Purpose Program for Assigning Sequence Reads to Genomic Features.” Bioinformatics 30 (7): 923–30.

Lindtner, Susan, Rinaldo Catta-Preta, Hua Tian, Linda Su-Feher, James D. Price, Diane E. Dickel, Vanille Greiner, et al. 2019. “Genomic Resolution of DLX-Orchestrated Transcriptional Circuits Driving Development of Forebrain GABAergic Neurons.” Cell Reports 28 (8): 2048–63.e8.

Lun, Aaron T. L., Davis J. McCarthy, and John C. Marioni. 2016. “A Step-by-Step Workflow for Low-Level Analysis of Single-Cell RNA-Seq Data with Bioconductor.” F1000Research 5 (August): 2122.

Lun, Aaron T. L., Samantha Riesenfeld, Tallulah Andrews, Dao, The Phuong, Tomas Gomes, and John C. Marioni. 2019. “EmptyDrops: Distinguishing Cells from Empty Droplets in Droplet-Based Single-Cell RNA Sequencing Data.” Genome Biology 20 (1): 1–9.

MacDonald, Adam, Brianna Lu, Maxime Caron, Nina Caporicci-Dinucci, Dale Hatrock, Kevin Petrecca, Guillaume Bourque, and Jo Anne Stratton. 2021. “Single Cell Transcriptomics of Ependymal Cells Across Age, Region and Species Reveals Cilia-Related and Metal Ion Regulatory Roles as Major Conserved Ependymal Cell Functions.” Frontiers in Cellular Neuroscience 15 (July): 703951.

Marsilje, Thomas H., Wei Pei, Bei Chen, Wenshuo Lu, Tetsuo Uno, Yunho Jin, Tao Jiang, et al. 2013. “Synthesis, Structure-Activity Relationships, and in Vivo Efficacy of the Novel Potent and Selective Anaplastic Lymphoma Kinase (ALK) Inhibitor 5-Chloro-N2-(2-Isopropoxy-5-Methyl-4-(piperidin-4-Yl)phenyl)-N4-(2-(isopropylsulfonyl)phenyl)pyrimidine-2,4-Diamine (LDK378) Currently in Phase 1 and Phase 2 Clinical Trials.” Journal of Medicinal Chemistry 56 (14): 5675–90.

Ma, Tong, Samuel Zheng Hao Wong, Bora Lee, Guo-Li Ming, and Hongjun Song. 2021. “Decoding Neuronal Composition and Ontogeny of Individual Hypothalamic Nuclei.” Neuron 109 (7): 1150–67.e6.

McInnes, Leland, John Healy, and James Melville. 2018. “UMAP: Uniform Manifold Approximation and Projection for Dimension Reduction.” http://arxiv.org/abs/1802.03426.

McMinn, J. E., C. W. Wilkinson, P. J. Havel, S. C. Woods, and M. W. Schwartz. 2000. “Effect of Intracerebroventricular Alpha-MSH on Food Intake, Adiposity, c-Fos Induction, and Neuropeptide Expression.” American Journal of Physiology. Regulatory, Integrative and Comparative Physiology 279 (2): R695–703.

Merkle, Florian T., Asif Maroof, Takafumi Wataya, Yoshiki Sasai, Lorenz Studer, Kevin Eggan, and Alexander F. Schier. 2015. “Generation of Neuropeptidergic Hypothalamic Neurons from Human Pluripotent Stem Cells.” Development 142 (4): 633–43.

Miranda-Angulo, Ana L., Mardi S. Byerly, Janny Mesa, Hong Wang, and Seth Blackshaw. 2014. “Rax Regulates Hypothalamic Tanycyte Differentiation and Barrier Function in Mice.” The Journal of Comparative Neurology 522 (4): 876–99.

Mok, Tony S. K., Lucio Crino, Enriqueta Felip, Ravi Salgia, Tommaso De Pas, Daniel S. W. Tan, and Laura Q. M. Chow. 2017. “The Accelerated Path of Ceritinib: Translating Pre-Clinical Development into Clinical Efficacy.” Cancer Treatment Reviews 55 (April): 181–89.

Morris, S. W., C. Naeve, P. Mathew, P. L. James, M. N. Kirstein, X. Cui, and D. P. Witte. 1997. “ALK, the Chromosome 2 Gene Locus Altered by the t(2;5) in Non-Hodgkin’s Lymphoma, Encodes a Novel Neural Receptor Tyrosine Kinase That Is Highly Related to Leukocyte Tyrosine Kinase (LTK).” Oncogene 14 (18): 2175–88.

Mountjoy, Kathleen G., Alexandre Caron, Kristina Hubbard, Avik Shome, Angus C. Grey, Bo Sun, Sarah Bould, et al. 2018. “Desacetyl-α-Melanocyte Stimulating Hormone and α-Melanocyte Stimulating Hormone Are Required to Regulate Energy Balance.” Molecular Metabolism 9 (March): 207–16.

Müller, Timo D., Matthias Blüher, Matthias H. Tschöp, and Richard D. DiMarchi. 2022. “Anti-Obesity Drug Discovery: Advances and Challenges.” Nature Reviews. Drug Discovery 21 (3): 201–23.

Nasser, Joseph, Drew T. Bergman, Charles P. Fulco, Philine Guckelberger, Benjamin R. Doughty, Tejal A. Patwardhan, Thouis R. Jones, et al. 2021. “Genome-Wide Enhancer Maps Link Risk Variants to Disease Genes.” Nature 593 (7858): 238–43.

Newman, Elizabeth A., Dong Won Kim, Jun Wan, Jie Wang, Jiang Qian, and Seth Blackshaw. 2018. “Foxd1 Is Required for Terminal Differentiation of Anterior Hypothalamic Neuronal Subtypes.” Developmental Biology 439 (2): 102–11.

Nicholas, Cory R., Jiadong Chen, Yunshuo Tang, Derek G. Southwell, Nadine Chalmers, Daniel Vogt, Christine M. Arnold, et al. 2013. “Functional Maturation of hPSC-Derived Forebrain Interneurons Requires an Extended Timeline and Mimics Human Neural Development.” Cell Stem Cell 12 (5): 573–86.

Nyberg, Solja T., G. David Batty, Jaana Pentti, Marianna Virtanen, Lars Alfredsson, Eleonor I. Fransson, Marcel Goldberg, et al. 2018. “Obesity and Loss of Disease-Free Years Owing to Major Non-Communicable Diseases: A Multicohort Study.” *The Lancet*. Public Health 3 (10): e490–97.

Okonechnikov, Konstantin, Ana Conesa, and Fernando García-Alcalde. 2015. “Qualimap 2: Advanced Multi-Sample Quality Control for High-Throughput Sequencing Data.” Bioinformatics 32 (2): 292–94.

Orthofer, Michael, Armand Valsesia, Reedik Mägi, Qiao-Ping Wang, Joanna Kaczanowska, Ivona Kozieradzki, Alexandra Leopoldi, et al. 2020. “Identification of ALK in Thinness.” Cell 181 (6): 1246–62.e22.

Palmer, Ruth H., Emma Vernersson, Caroline Grabbe, and Bengt Hallberg. 2009. “Anaplastic Lymphoma Kinase: Signalling in Development and Disease.” Biochemical Journal 420 (3): 345–61.

Pantazis, Caroline B., Andrian Yang, Erika Lara, Justin A. McDonough, Cornelis Blauwendraat, Lirong Peng, Hideyuki Oguro, et al. 2022. “A Reference Human Induced Pluripotent Stem Cell Line for Large-Scale Collaborative Studies.” Cell Stem Cell 29 (12): 1685–1702.e22.

Pietzner, Maik, Eleanor Wheeler, Julia Carrasco-Zanini, Adrian Cortes, Mine Koprulu, Maria A. Wörheide, Erin Oerton, et al. 2021. “Mapping the Proteo-Genomic Convergence of Human Diseases.” Science 374 (6569): eabj1541.

Pons, Pascal, and Matthieu Latapy. 2005. “Computing Communities in Large Networks Using Random Walks (long Version).” http://arxiv.org/abs/physics/0512106.

Prevot, Vincent, Bénédicte Dehouck, Ariane Sharif, Philippe Ciofi, Paolo Giacobini, and Jerome Clasadonte. 2018. “The Versatile Tanycyte: A Hypothalamic Integrator of Reproduction and Energy Metabolism.” Endocrine Reviews 39 (3): 333–68.

Qiao, Liping, Bonggi Lee, Brice Kinney, Hyung Sun Yoo, and Jianhua Shao. 2011. “Energy Intake and Adiponectin Gene Expression.” American Journal of Physiology. Endocrinology and Metabolism 300 (5): E809–16.

Qi, Ting, Yang Wu, Jian Zeng, Futao Zhang, Angli Xue, Longda Jiang, Zhihong Zhu, et al. 2018. “Identifying Gene Targets for Brain-Related Traits Using Transcriptomic and Methylomic Data from Blood.” Nature Communications 9 (1): 2282.

Rajamani, Uthra, Andrew R. Gross, Brooke E. Hjelm, Adolfo Sequeira, Marquis P. Vawter, Jie Tang, Vineela Gangalapudi, et al. 2018. “Super-Obese Patient-Derived iPSC Hypothalamic Neurons Exhibit Obesogenic Signatures and Hormone Responses.” Cell Stem Cell 22 (5): 698–712.e9.

Raudvere, Uku, Liis Kolberg, Ivan Kuzmin, Tambet Arak, Priit Adler, Hedi Peterson, and Jaak Vilo. 2019. “g:Profiler: A Web Server for Functional Enrichment Analysis and Conversions of Gene Lists (2019 Update).” Nucleic Acids Research 47 (W1): W191–98.

Reshetnyak, Andrey V., Paolo Rossi, Alexander G. Myasnikov, Munia Sowaileh, Jyotidarsini Mohanty, Amanda Nourse, Darcie J. Miller, Irit Lax, Joseph Schlessinger, and Charalampos G. Kalodimos. 2021. “Mechanism for the Activation of the Anaplastic Lymphoma Kinase Receptor.” Nature 600 (7887): 153–57.

Roberts, Brandon L., Camdin M. Bennett, Julie M. Carroll, Sarah R. Lindsley, and Paul Kievit. 2019. “Early Overnutrition Alters Synaptic Signaling and Induces Leptin Resistance in Arcuate Proopiomelanocortin Neurons.” Physiology & Behavior 206 (July): 166–74.

Santos, David P., Evangelos Kiskinis, Kevin Eggan, and Florian T. Merkle. 2016. “Comprehensive Protocols for CRISPR/Cas9-Based Gene Editing in Human Pluripotent Stem Cells.” Current Protocols in Stem Cell Biology 38 (August): 5B.6.1–5B.6.60.

Secher, Anna, Jacob Jelsing, Arian F. Baquero, Jacob Hecksher-Sørensen, Michael A. Cowley, Louise S. Dalbøge, Gitte Hansen, et al. 2014. “The Arcuate Nucleus Mediates GLP-1 Receptor Agonist Liraglutide-Dependent Weight Loss.” The Journal of Clinical Investigation 124 (10): 4473–88.

Shaw, A. T., R. Mehra, D. S. W. Tan, E. Felip, L. Q. Chow, D. R. Camidge, J. F. Vansteenkiste, et al. 2014. “Evaluation of Ceritinib-Treated Patients (Pts) with Anaplastic Lymphoma Kinase Rearranged (Alk+) Non-Small Cell Lung Cancer (Nsclc) and Brain Metastases in the Ascend-1 Study.” Annals of Oncology: Official Journal of the European Society for Medical Oncology / ESMO 25 (September): iv455.

Shimogori, Tomomi, Daniel A. Lee, Ana Miranda-Angulo, Yanqin Yang, Hong Wang, Lizhi Jiang, Aya C. Yoshida, et al. 2010. “A Genomic Atlas of Mouse Hypothalamic Development.” Nature Neuroscience 13 (6): 767–75.

Siletti, Kimberly, Rebecca Hodge, Alejandro Mossi Albiach, Lijuan Hu, Ka Wai Lee, Peter Lönnerberg, Trygve Bakken, et al. 2022. “Transcriptomic Diversity of Cell Types across the Adult Human Brain.” bioRxiv. https://doi.org/10.1101/2022.10.12.511898.

Soda, Manabu, Young Lim Choi, Munehiro Enomoto, Shuji Takada, Yoshihiro Yamashita, Shunpei Ishikawa, Shin-Ichiro Fujiwara, et al. 2007. “Identification of the Transforming EML4-ALK Fusion Gene in Non-Small-Cell Lung Cancer.” Nature 448 (7153): 561–66.

Ste Marie, L., G. I. Miura, D. J. Marsh, K. Yagaloff, and R. D. Palmiter. 2000. “A Metabolic Defect Promotes Obesity in Mice Lacking Melanocortin-4 Receptors.” Proceedings of the National Academy of Sciences of the United States of America 97 (22): 12339–44.

Steuernagel, Lukas, Brian Y. H. Lam, Paul Klemm, Georgina K. C. Dowsett, Corinna A. Bauder, John A. Tadross, Tamara Sotelo Hitschfeld, et al. 2022. “HypoMap—a Unified Single-Cell Gene Expression Atlas of the Murine Hypothalamus.” Nature Metabolism 4 (10): 1402–19.

Sullivan, Andrew I., Matthew J. Potthoff, and Kyle H. Flippo. 2022. “Tany-Seq: Integrated Analysis of the Mouse Tanycyte Transcriptome.” Cells 11 (9). https://doi.org/10.3390/cells11091565.

Sun, Feng, Sanbao Chai, Lishi Li, Kai Yu, Zhirong Yang, Shanshan Wu, Yuan Zhang, Linong Ji, and Siyan Zhan. 2015. “Effects of Glucagon-like Peptide-1 Receptor Agonists on Weight Loss in Patients with Type 2 Diabetes: A Systematic Review and Network Meta-Analysis.” Journal of Diabetes Research 2015 (January): 157201.

Sun, Zhiqi, Jingyi Gong, Han Wu, Wenyi Xu, Lizhen Wu, Dijin Xu, Jinlan Gao, et al. 2013. “Perilipin1 Promotes Unilocular Lipid Droplet Formation through the Activation of Fsp27 in Adipocytes.” Nature Communications 4: 1594.

Üner, Aykut Göktürk, Onur Keçik, Paula G. F. Quaresma, Thiago M. De Araujo, Hyon Lee, Wenjing Li, Hyun Jeong Kim, Michelle Chung, Christian Bjørbæk, and Young-Bum Kim. 2019. “Role of POMC and AgRP Neuronal Activities on Glycaemia in Mice.” Scientific Reports 9 (1): 13068.

Vernersson, Emma, Nelson K. S. Khoo, Maria L. Henriksson, Göran Roos, Ruth H. Palmer, and Bengt Hallberg. 2006. “Characterization of the Expression of the ALK Receptor Tyrosine Kinase in Mice.” Gene Expression Patterns: GEP 6 (5): 448–61.

Võsa, Urmo, Annique Claringbould, Harm-Jan Westra, Marc Jan Bonder, Patrick Deelen, Biao Zeng, Holger Kirsten, et al. 2021. “Large-Scale Cis- and Trans-eQTL Analyses Identify Thousands of Genetic Loci and Polygenic Scores That Regulate Blood Gene Expression.” Nature Genetics 53 (9): 1300–1310.

Wang, Liheng, Alain J. De Solis, Yossef Goffer, Kathryn E. Birkenbach, Staci E. Engle, Ross Tanis, Jacob M. Levenson, et al. 2019. “Ciliary Gene RPGRIP1L Is Required for Hypothalamic Arcuate Neuron Development.” JCI Insight 4 (3). https://doi.org/10.1172/jci.insight.123337.

Wang, Liheng, Kana Meece, Damian J. Williams, Kinyui Alice Lo, Matthew Zimmer, Garrett Heinrich, Jayne Martin Carli, et al. 2015. “Differentiation of Hypothalamic-like Neurons from Human Pluripotent Stem Cells.” The Journal of Clinical Investigation 125 (2): 796–808.

Wang, Weidong, J. Fredrik Grimmer, Thomas R. Van De Water, and Thomas Lufkin. 2004. “Hmx2 and Hmx3 Homeobox Genes Direct Development of the Murine Inner Ear and Hypothalamus and Can Be Functionally Replaced by Drosophila Hmx.” Developmental Cell 7 (3): 439–53.

Weeks, Elle M., Jacob C. Ulirsch, Nathan Y. Cheng, Brian L. Trippe, Rebecca S. Fine, Jenkai Miao, Tejal A. Patwardhan, et al. 2020. “Leveraging Polygenic Enrichments of Gene Features to Predict Genes Underlying Complex Traits and Diseases.” medRxiv. https://doi.org/10.1101/2020.09.08.20190561.

Woodling, Nathaniel S., Benjamin Aleyakpo, Miranda Claire Dyson, Lucy J. Minkley, Arjunan Rajasingam, Adam J. Dobson, Kristie H. C. Leung, et al. 2020. “The Neuronal Receptor Tyrosine Kinase Alk Is a Target for Longevity.” Aging Cell 19 (5): e13137.

Yang, Jian, S. Hong Lee, Michael E. Goddard, and Peter M. Visscher. 2011. “GCTA: A Tool for Genome-Wide Complex Trait Analysis.” American Journal of Human Genetics 88 (1): 76–82.

Yengo, Loic, Julia Sidorenko, Kathryn E. Kemper, Zhili Zheng, Andrew R. Wood, Michael N. Weedon, Timothy M. Frayling, et al. 2018. “Meta-Analysis of Genome-Wide Association Studies for Height and Body Mass Index in ∼700000 Individuals of European Ancestry.” Human Molecular Genetics 27 (20): 3641–49.

Yeo, Giles S. H., Daniela Herrera Moro Chao, Anna-Maria Siegert, Zoe M. Koerperich, Mark D. Ericson, Stephanie E. Simonds, Courtney M. Larson, et al. 2021. “The Melanocortin Pathway and Energy Homeostasis: From Discovery to Obesity Therapy.” Molecular Metabolism 48 (June): 101206.

Yeo, Giles S. H., Emma J. Lank, I. Sadaf Farooqi, Julia Keogh, Benjamin G. Challis, and Stephen O’Rahilly. 2003. “Mutations in the Human Melanocortin-4 Receptor Gene Associated with Severe Familial Obesity Disrupts Receptor Function through Multiple Molecular Mechanisms.” Human Molecular Genetics 12 (5): 561–74.

Zamo, Alberto, Roberto Chiarle, Roberto Piva, Jennifer Howes, Yan Fan, Marco Chilosi, David E. Levy, and Giorgio Inghirami. 2002. “Anaplastic Lymphoma Kinase (ALK) Activates Stat3 and Protects Hematopoietic Cells from Cell Death.” Oncogene 21 (7): 1038–47.

Zhan, Cheng, Jingfeng Zhou, Qiru Feng, Ju-En Zhang, Shuailiang Lin, Junhong Bao, Ping Wu, and Minmin Luo. 2013. “Acute and Long-Term Suppression of Feeding Behavior by POMC Neurons in the Brainstem and Hypothalamus, Respectively.” The Journal of Neuroscience: The Official Journal of the Society for Neuroscience 33 (8): 3624–32.

Zhang, Shiqi, Guowen Liu, Chuang Xu, Lei Liu, Qiang Zhang, Qiushi Xu, Hongdou Jia, Xiaobing Li, and Xinwei Li. 2018. “Perilipin 1 Mediates Lipid Metabolism Homeostasis and Inhibits Inflammatory Cytokine Synthesis in Bovine Adipocytes.” Frontiers in Immunology 9 (March): 467.

Zhu, Zhihong, Futao Zhang, Han Hu, Andrew Bakshi, Matthew R. Robinson, Joseph E. Powell, Grant W. Montgomery, et al. 2016. “Integration of Summary Data from GWAS and eQTL Studies Predicts Complex Trait Gene Targets.” Nature Genetics 48 (5): 481–87.

Zupančič, Maja, Evgenii Tretiakov, Zoltán Máté, Ferenc Erdélyi, Gábor Szabó, Frédéric Clotman, Tomas Hökfelt, Tibor Harkany, and Erik Keimpema. 2023. “Brain-Wide Mapping of Efferent Projections of Glutamatergic (Onecut3) Neurons in the Lateral Mouse Hypothalamus.” Acta Physiologica 238 (3): e13973.

